# Systemic inflammation causes microglial dysfunction with a mixed AD-like pathology

**DOI:** 10.1101/2020.07.27.223198

**Authors:** Praveen Bathini, Isabel Dupanloup, Elena Zenaro, Eleonora Terrabuio, Amrei Fischer, Edona Ballabani, Marie-Agnes Doucey, Lavinia Alberi

## Abstract

**Background:** Alzheimer’s disease (AD) is the primary cause of cognitive deficit in elderly humans. Late-onset AD (LOAD) is sporadic, multifactorial, non-Mendelian accounting at present for 95% of the cases in contrast to the genetic form. Risk factors for sporadic AD include Gene: Environment interactions. There is increasing evidence that lifestyle and environmental stress such as infection and chronic inflammation are underlying culprits of neurodegenerative dementia. Dementias that share or mimic pathological processes of AD include cerebrovascular diseases, Lewy body disease, TDP-43 proteinopathy. To date, very few mouse models reproduce the pathophysiological progression of mixed-vascular-AD, while the majority of studies have employed transgenic animals reproducing the familial form.

**Methods:** We have re-engineered the Polyinosinic:polycytidylic acid (PolyI:C) sterile infection model in wildtype C57Bl6 mice to obtain chronic low-grade systemic inflammation. We have conducted a cross-sectional analysis of aging PolyI:C and Saline control mice (3 months, 6 months, 9 months and 16 months), taking the hippocampus as a reference brain region, based on its vulnerability, and compared the brain aging phenotype to AD progression in humans with mild AD, severe AD and Controls (CTL), parallely in Vascular dementia (VaD) patient specimens.

**Results:** We found that PolyI:C mice display both peripheral and central inflammation with a peak at 6 months, associated with memory deficits. The hippocampus is characterized by a pronounced and progressive tauopathy. In PolyI:C brains, microglia undergo aging-dependent morphological rearrangements progressively adopting a phagocytic phenotype. Transcriptomic analysis reveals a profound change in gene expression over the course of aging, with a peak in differential expression at 9 months. We confirm that the proinflammatory marker *Lcn2* is one of the genes with the strongest upregulation in PolyI:C mice upon aging. Validation in brains from patients with increasing severity of AD and VaD shows a reproducibility of some gene targets in vascular dementia specimens rather than AD ones, in which only GFAP is strongly increased at the severe stages.

**Conclusions:** The PolyI:C model of sterile infection demonstrates that peripheral chronic inflammation is sufficient to cause neuropathological processes resembling a mixed-VaD-AD phenotype, with progressive tau hyperphosphorylation, changes in microglia morphology, astrogliosis and gene reprogramming reflecting increased neuroinflammation, vascular remodeling and the loss of neuronal functionality seen to some extent in humans.

## Background

Alzheimer’s disease (AD) is the most common form of dementia, with vascular dementia (VaD) accounting for the second most common form. AD is a progressive neurodegenerative disorder characterized by amyloid-*β* plaques and neurofibrillary tangles resulting in cognitive decline and neuronal loss. Most of the AD cases are familial and account for 5% of the cases whereas 95% cases are sporadic with a late-onset AD (Alonso, Grundke-Iqbal, and Iqbal 1996; Masters et al. 2015). While the familial AD (FAD) has been linked mostly to the mutations in the amyloid precursor protein (APP) and presenilin-1 &2 genes (PS1 and PS2), the etiology of late-onset or sporadic AD affecting people over 65 years of age is still not clearly understood. An increasing body of literature shows that mixed pathologies involving AD with cerebrovascular diseases, or Lewy body disease, or TDP-43 proteinopathy (10-78% prevalence) (Rahimi and Kovacs 2014) with cerebrovascular and sporadic AD (over 20% of cases) representing the most common mixed pathology in advanced aging (Toledo et al. 2013; Lin et al. 2019; Santos et al. 2017). Scientific studies show the involvement of both genetic and environmental factors in sporadic AD. Some of these genes include ABCA7, APOE, BIN1, CD2AP, CD33, CLU, CR1, EPHA1, MS4A4A/MS4A4E/MS4A6E, PICALM, and SORL1 (Barber 2012), of which a large proportion are immunological factors. .Modifiable risk factors for AD include inflammation, viral infections, Type 2 diabetes, vascular disorders, and hypertension (Edwards et al. 2019; Stozická, Zilka, and Novák 2007). Among these risk factors, inflammation is considered as a central mechanism in AD and infectious agents like bacteria and viruses might play a critical role in sustaining chronic inflammation and inducing neuroinflammation in particular in subjects carrying the aforementioned genetic variants. SORL1 and APOE have been also associated with Cerebral Amyloid Angiopathy (CAA) (Du et al. 2019) supporting vascular inflammation as comorbidity in AD. Notably, Cerebral Amyloid Angiopathy (CAA) is associated with higher CERAD scores, higher phase of Aβ deposition, and higher Braak stages (Thal et al. 2002). Since the last decade, many epidemiological studies in humans have proposed the link between dysfunction of the immune system and AD pathogenesis (Morgan et al. 2019; Engelhart et al. 2004; Leung et al. 2013), yet much less attention has been given to mixed pathologies. Earlier studies indicate that peripheral inflammation can prompt central immune responses (Perry, Cunningham, and Holmes 2007) and an immune memory persisting in the brain (Saeed et al. 2014) causing neurodegeneration. Also, abnormal expression of these immune factors in response to stress or infection during pregnancy can lead to neurodevelopmental disorders and behavioral deficits in the offspring (Bilbo et al. 2018) and may underlie neurodegenerative processes later in life (Knuesel et al. 2014).

Here, we investigated pathophysiological response using a wild type mouse model exposed to viral-like immune activation, using Polyinosinic:polycytidylic acid (PolyI:C), in late pregnancy and the young offspring. The second immune challenge bolus was designed to achieve sustained chronic inflammation deviating from the original model (Krstic et al. 2012) that could not be replicated by other studies (Giovanoli et al. 2015). The new optimized PolyI:C mouse model reveals long term systemic & central immune response resulting in significant proteinopathy associated with spatial memory deficit, microglia remodeling, transcriptomic changes leading to increased lipid metabolism and vascular factors activation, resembling a mixed vascular-AD pathophysiology.

## Materials and Methods

### Animals

Wildtype C57BL/6J mice were used for the study. Animal experimentation was approved by the animal experiment committee, University of Fribourg (Protocol no. 2016_32_FR registered 01/01/2017). The animals were fed ad libitum and housed in a room equipped with automatically controlled temperature (21-25 °C), humidity (50%), and with a 12 hrs light/ dark cycle.

### Human tissue

Frozen Human tissue samples from the entorhinal cortex including the hippocampal area were procured from the Medical Research Council Brain Bank for Dementia Research, Oxford, UK, and Stanford brain bank. We received frozen hippocampal brain tissue samples from 9 controls, 5 moderate, and 10 Severe sporadic AD patients (Suppl. Table 1). The use of human tissue has been approved by the Ethical Commission of the Brain Bank for Dementia UK (OBB443 registered 1/05/2017 and OB344 registered 1/02/2014), Stanford (Stanford IRB), and the Ethical Commission from the Canton of Fribourg and Vaud (N. 325/14). Vascular dementia hippocampal specimens and healthy age-matched controls were obtained from the Netherlands Brain Bank (NBB. 1251). All experiments conducted on human tissue comply with the WMA Declaration of Helsinki.

### Treatment

To investigate the early postnatal treatment and microglial priming the systemic Poly I:C challenges were optimized from the already existing model with PolyI:C (Krstic et al. 2012). Female C57BL/6J mice 6-8 weeks old were housed together with the males for mating. Vaginal plugs were assessed and pregnant mice with gestation day (GD) 17 were injected intravenously (i.v.) with 5mg/kg PolyI:C (Polyinosinic-polycytidylic acid; P9582, Sigma, USA). Aliquots of 5mg/ml were prepared by resuspending in the sterile 0.9% saline and were stored at −20°C until further use. For control experiments sterile 0.9% saline was used. To mimic the chronic inflammatory conditions like in humans, prenatally challenged offspring were given a second immune challenge with an intraperitoneal (i.p.) PolyI:C injection at 20mg/kg dose or sterile saline for the control experimental mice (Fig. 1A).

**Figure 1.**
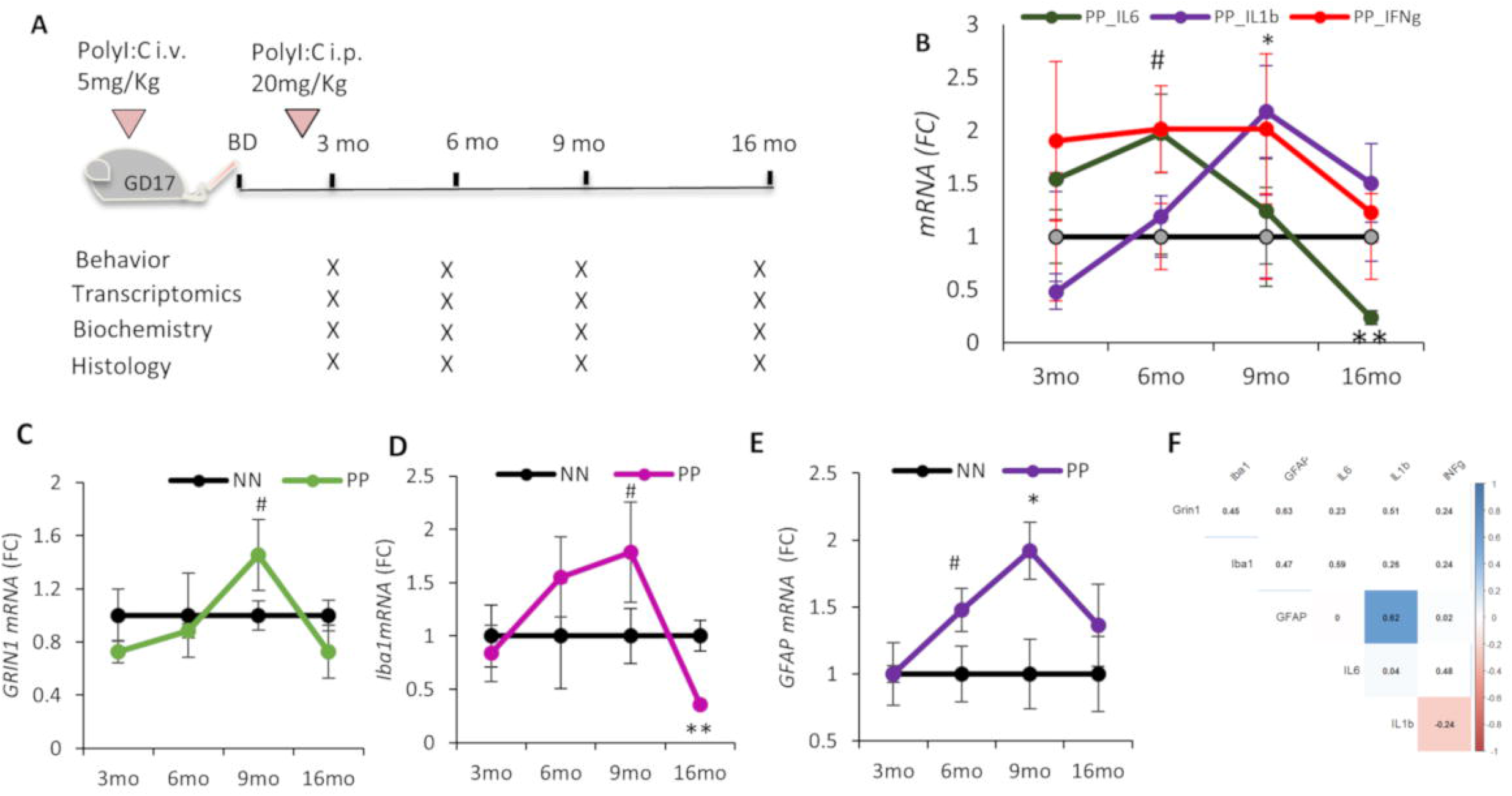
The neuroinflammatory evolution in aging PolyI:C mice. **(A)** Timeline of the experimental procedure and PolyI:C injections. **(B)** RT-PCR analysis of proinflammatory cytokines in the hippocampus after the PolyI:C treatment. Transcript analysis for **(C)** neuronal (GRIN1), **(D)** microglia (Iba1), and **(E)** astroglial (GFAP) markers. **(F)** The correlation matrix indicates the associations between time series of Grin1, Iba1, GFAP with the examined cytokines. Values represented as mean ± SEM, n=4-5 mice per age, and treatment. GD=gestational day, BD=birthday. *p<0.05, **p<0.005, #p=0.07. Student’s t-test.

### Behavioural experiments

Anxiety was tested at 3, 6, 9 and 16 months of age in PP and NN mice using the Open field test registering the time spent in the center, while elevated O-maze was used at 3 to 6 months only recording both the time spent in the open unprotected arm and time spent in the light. Working memory was tested using the Y-maze alternation task and expressed as % of alternation and number of arm entries during a 5 minutes exploration-window according to previously published protocols (Crawley and Bailey 2008; Miedel et al. 2017). Mice during the behavior were videotaped and Videos were analyzed using Image J-built in macros for automated video-tracking (Supplementary Material and Methods) (Brai and Alberi 2015).

### Tissue Processing

Animals were first deeply anesthetized with pentobarbital sodium (100mg/kg, i.p), later toe pinch pain reflex was checked. After the abdominal opening, mice were transcardially perfused with 0.9% sterile saline, brains were harvested and cut into two hemispheres. One hemisphere was dissected to collect the hippocampus and further dissected in ice-cold saline solution to obtain the Corpus Ammonis (CA) fields, removing the dentate gyrus (Brai et al. 2015). The tissue samples were collected into eppendorfs and were flash-frozen in liquid nitrogen and stored at −80°C until further use. Another hemisphere was post-fixed in 4% PFA for 1 day, followed by immersion in 30% sucrose at 4°C sucrose and then embedded in an OCT block for cryosectioning at 35μm thickness (Leica, Germany) and used for histological studies.

### Plasma collection

The whole blood collected via cardiac puncture was transferred into the EDTA-coated eppendorfs. Cells were separated from the plasma by centrifugation for 15 min at 2000 × g at 4°C before extracting the plasma. The plasma samples were then stored at −80°C until further use.

### Isolation of leukocytes from mouse tissues and Flow cytometry

#### Blood

Mice were anesthetized via isoflurane inhalation. Blood samples were collected through a retro-orbital not-surgical procedure by sodium heparinized capillaries. Dextran 1% and Heparin 10 U/ml were added to blood in a 1:1 ratio. After erythrocytes sedimentation (1 hour), the overlying supernatant plasma-dextran suspension of leukocytes was washed in 1X PBS. Red blood cell lysis was performed adding 3 ml of NaCl 0.2% for 40 seconds and then 7 ml of NaCl 1.2%. Next, cells were washed and stained.

#### Brain

Mice were anesthetized and quickly perfused through the left cardiac ventricle by injection of cold PBS 1X Ca_2_Mg_2_ 1 mM. After meninges removal, brains were collected. Mechanical digestion through gentleMACS™ Octo Dissociator (Miltenyi Biotec) and subsequent enzymatic digestion with 2 mg/ml of collagenase (C0130; Sigma-Aldrich, USA) and 40 U/ml of DNase (EN0521; Invitrogen, Life Technologies, USA) at 37°C for 45 minutes in water bath were performed. Cells were isolated by passing the digested tissue through a 70-mm cell strainer, resuspended in 30% Percoll (GE Healthcare, USA), and loaded onto 70% Percoll (GE Healthcare, USA). After centrifugation at 2500 rpm for 20 minutes at 4°C, cells were removed from interphase, washed, and stained.

Cells were treated with an antibody against Fc receptor [anti-mouse CD16/32, Clone: 2.4G2; Becton Dickinson (BD), USA] and labelled with BD Horizon Brilliant™ Stain Buffer (BD, USA) to improve staining quality and with 2,5 mL per sample of each of the following anti-mouse mAbs: Ly6G BB515 (Clone: 1A8; BD, USA), CD45 BV786 (Clone, 104 RUO; BD, USA), CD11b BV421 (Clone: R1-2; BD, USA), F4/80 (Clone: T45-2342 RUO; BD, USA), CD11c BV 605 (Clone: HL3; BD, USA), Siglec F PE (Clone: S17007L; BD, USA),B220 APC (Clone: RA3-6B2 RUO; BD, USA). Cells were stained for 15 minutes at 4°C in the dark according to manufacturer’s instructions. After washing, cells were incubated with 7AAD (BioLegend, USA) fluorescent intercalator for 5 minutes at 4°C in the dark. Samples were acquired through LSRFortessa X-20 (BD, USA). Data were analyzed through FlowJo™ Software. In particular, after doublets removal, 7AAD^−^ alive cells were selected. A specific gating strategy was used to identify subpopulations of leukocytes in blood and brain samples. The following CD45^+^ cell populations were detected and analyzed: Ly6G^+^ CD11b^+^ B220^−^ Neutrophils, Siglec F^+^ CD11b^−^ CD11c^−^ Eosinophils, F4/80^+^ CD11b^+^ SiglecF^−^ Monocytes. Using the DownSample Plugin developed by FlowJo™ (https://docs.flowjo.com/seqgeq/dimensionality-reduction/downsample/), the number of CD45^+^ events in data matrix was reduced to a maximum of 30.000.

### Bead-based Immunoassay

To study the PolyI:C effect on plasma inflammatory profile, we measured chemokines IL-6, IL-10, MCP-1, and TNF-α using the premixed inflammation panel (PN: C282251A, Aimplex Biosciences, USA) according to the manufacturer’s instructions. Briefly, 45 μl of the capture bead working solution, 30 μl of assay buffer, and 15 μl of the sample were incubated on the shaker (700 rpm) for 60 minutes at room temperature. After 3 washes with 100 μl washing buffer, 25 μl of biotinylated antibody working solution was added and incubated on the shaker for 30 min at room temperature with protection from light. After 3 washes, 25 μL of streptavidin-phycoerythrin working solution was added, shaken for 20 minutes in the dark. Finally, the beads were resuspended on 150 μl of 1x reading buffer and read on Flow Cytometry Analyzer (LSR-IIa; BD, USA) acquiring about 50 events per chemokine. The FCS files were analyzed on Flowing software (http://flowingsoftware.btk.fi/) and the inflammatory levels were determined based on 5-parameter logistic curve fitting.

### Immunoassay of Amyloid-β_1-42_

Frozen cortical tissue samples were homogenized with lysis buffer containing 20 mM Tris-HCl (pH 7.5), 150 mM NaCl, 1 mM disodium EDTA, 1 mM EGTA, 1% Triton, 2.5 mM sodium pyrophosphate, 1 mM β-glycerophosphate (G9422; Sigma, Germany), phosphatases inhibitor such as Sodium Orthovanadate (1mM, 450243; Sigma, Germany) and proteases inhibitor cocktail 1:100 (3749.1, Roth, Germany) and the lysates were stored at −80°C until further analysis. The sample and reagents preparation has been carried out according to the Milliplex Map kit Human Amyloid and Tau Magnetic Bead panel (Cat.# HNABTMAG-68K) manufacturer’s protocol. Briefly, cortical lysates were diluted 1:2 with assay buffer. Standards, controls and samples were added to the appropriate wells of the 96 well filter plate. Biotinylated detection antibodies and magnetic mixed antibody-immobilized beads were added, sealed with aluminum foil, and incubated overnight in the dark at room temperature on a shaker (700rpm). The next day following washes, Streptavidin-phycoerythrin was added and incubated in the dark for 30 minutes at room temperature on a shaker (700rpm). Finally, the beads were suspended in 100 μl Sheath fluid and transferred to the 96 well flat bottom plate to be read on the FlexMap 3D system equipped with xPONENT software (Merck, Germany). The median fluorescence intensity (MFI) was analyzed using a 5-parameter logistic curve-fitting method and analyte concentrations were calculated.

### Nucleic acid and protein extraction

Fresh frozen hippocampal tissue specimens from mice and humans are stored at −80°C until utilized for RNA extraction using peqGOLD TriFast™ (peqGOLD, Germany) according to the manufacturer’s instructions to obtain RNA and proteins. The RNA pellet was extracted from the aqueous phase, air-dried, resuspended in 25μl of nuclease-free water (Promega). The concentration of the RNA samples was measured using the Qubit 3.0 Fluorometer (High sensitivity, Invitrogen) and stored at −80°C until further use. Proteins were extracted from the organic phase, cleaned with Ethanol, and the dried pellet resuspended with a 150 μl buffer containing 8M Urea in 4% (w/v) CHAPS and protease inhibitor (1:100 Carl Roth, Germany). The protein concentrations were determined using the Bradford assay method (Roth, Germany) and later the samples were stored until use at −80°C.

### RNA library preparation and sequencing

Synthesis and amplification of cDNA were performed as per the Illumina TruSeq Stranded mRNA protocol. RNA extracted from the hippocampal specimens from all the age groups (3m, 6m, 9m, 16m) was used for this transcriptomics study. RNA integrity was determined with the Fragment Analyzer 5200 (Agilent). Samples with RNA integrity number (RIN) > 8 were used for the experiment. An input material of 1 μg of total RNA from each sample was used for library preparation with Illumina TruSeq Stranded mRNA kit (Cat: 20020595). Illumina TrueSeq Combinatorial dual (CD) indexes were used during the ligation, DNA fragments enriched using the PCR to amplify the amount of DNA in the library. The quality of the libraries is determined using the Standard High sensitivity NGS Fragment analysis kit (DNF-474, 1-6000 base pair) on the Agilent Fragment analyzer (Agilent, USA), yielding approximately 260 bp size fragments. The cDNA libraries were pooled in equivalent amounts. The libraries were denatured and diluted using standard library quantification and quality control procedures recommended as per the NextSeqprotocol. For a sequencing control, PhiX library was prepared and combined with the pooled prepared libraries. A final concentration of 1.5 pM library was sequenced on Illumina NextSeq 500 system (High output 150 cycles) to generate 20 million of 2 ×75 bp pair-end reads per library.

### Bioinformatics Analysis

In brief, hippocampal paired-end libraries from all the age groups (3m, 6m, 9m, 16m) in the saline treated and PolyI:C treated mice (n=3 each) were sequenced for the study. To generate the differentially expressed transcripts from the RNA sequencing data the following bioinformatics analysis pipeline was performed. 1) Remove adapter sequences and remove low quality (flanking N) bases from each read using cutadapt version 2.3; 2) Alignment of the reads to the reference genome using RNA-seq aligner STAR version 2.6; 3) Get basic alignment stats with RSeQC version 2.6.4; 4) Get read distribution with RSeQC version 2.6.4; 5) Get gene body coverage with RSeQC version 2.6.4; 6) Get read counts for genes with htseq-count release_0.11.1; 7) perform differential expression analysis with DESEQ2.

Further functional enrichments were performed using the ClueGO (Bindea et al. 2009) which integrates Gene ontology (GO), KEGG (Kyoto Encyclopedia of Genes and Genomes), Wikipathway, Reactome pathway analysis creating a functional pathway network.

### Reverse transcription-polymerase chain reaction (RT-PCR)

1μg of total RNA from the human and mouse brain specimen was reverse-transcribed using the M-MLV reverse transcriptase (Promega) for efficient synthesis of the first-strand cDNA. Gene expression analysis was done by RT-PCR (GoTaq qPCR Master Mix, Promega, USA) using gene specific primers (Suppl. Table 2) on Mic qPCR Cycler (BioMolecular Systems, USA). Expression levels of genes of interest were determined using the ΔΔCT method; the levels of the mRNAs of interest were normalized against the levels of the housekeeping gene, β-actin.

### Antibodies & reagents

Following antibodies were used for western blot immunolabeling, immunohistochemistry experiments in this study:

Rabbit anti pTau 231 (1:500, ab151559, Abcam, UK), Rabbit anti pTau 205 (1:500, ab181206, Abcam, UK), goat anti-tau (1.25 μg/mL; AF3494, R&D systems, USA), mouse anti-β actin, 1:2000 (sc-81178; Santa Cruz Biotechnology, USA), Rabbit anti GFAP (1:500, ab68428, Abcam, UK), Goat anti GFAP (1:500, SAB2500462, Sigma), Goat anti Iba1 (1:500, ab5076, Abcam, UK), Mouse anti-amyloid beta 6E10 clone (1:500, cat:803014; Biolegend, USA), rabbit anti-synaptophysin (1:500; ab14692, Abcam, UK), Rabbit anti-amyloid beta 42 (Aβ-42; 1:500; D54D2 XP, Cell Signaling, USA), Rabbit anti-Lipocalin-2/LCN2 (1:500, 50060-RP02; Sino Biological, China), mouse anti-NF200 (1:500, cat:1178709; Boehringer Mannheim Biochemica, Germany), Nile Red (2 μg/mL, N3013, Sigma, USA), Thioflavin-S (100 mM; CAS 1326-12-1, Santa Cruz Biotechnology, USA)

The secondary antibodies used in the study for immunolabeling were: directly conjugated to Cy2, Cy3, or Cy5 in donkey raised secondary antibodies (all 1:1000; Jackson Immuno Europe, UK). After the completion of immunofluorescence protocols, the sections were stained with DAPI to visualize nuclear morphology, mounted on Super frost slides (Thermofisher, USA), and coverslipped with a custom made Polyvinyl alcohol (PVA) and 1,4-diazabicyclo[2.2.2]octane (DABCO)-based mounting media.

### Western blot

Hippocampal lysates supplemented were denatured at 95°C using the 2-Mercaptoethanol based loading buffer. Proteins were separated using the SDS-polyacrylamide gel electrophoresis and western blot procedure. Custom made 8-10% gels were used to run the samples for 1.5 hours at 110 voltage. The proteins were then transferred to a precut 0.2 μm nitrocellulose membrane (Cat: 1620146; Biorad, USA) using a wet transfer method (Biorad). Later the membranes were incubated in a blocking solution containing 0.5% bovine serum albumin (Art. No. 8076.4, Roth, Germany), 1xTBS, and 0.1%Tween-20 for 30 minutes at room temperature on a rotating shaker. The membranes were then incubated with primary antibodies overnight at 4°C with gentle agitation. On the second day, the membranes were rinsed with TBS-Tween solution 3 × 5 minutes, followed by a 1-hour incubation with the fluorescently conjugated secondary antibodies, rinsed again and air-dried covered by aluminum foil. The proteins were detected using the Omega Lum (Labgene, CH). The optical density of the protein bands was determined using Image J software and normalized to the Beta-actin control. For an accurate total protein quantification REVERT stain (LI-COR Biosciences-GmbH, Germany) was also employed.

### Fluorescent immunohistochemistry

The mice sagittal tissue sections from the anti-freezing medium were taken and then mounted onto the superfrost glass slides, air-dried for 1 hour, and washed twice in Distilled water 5 minutes each. To access epitopes slides were incubated at 65°C (water bath) for 20 minutes in 10mM Sodium citrate (pH 6) containing 0.05% Tween (Preheat buffer). Thereafter, sections were washed 3 times 5 minutes each in 1x Trizma-based salt solution (TBS), at room temperature, and once in a TBST buffer (1x TBS containing 0.1% Triton solution). Sections were blocked in the blocking solution (TBST buffer containing 10% fetal bovine serum) for 1 hour at room temperature in a humid chamber. Further, the sections were incubated with the primary antibody in the TBST buffer containing 1% fetal bovine serum at 4°C in the refrigerator overnight. The next day, sections were washed 3 times for 5 minutes each in the TBS buffer before being incubated with the fluorescently labeled secondary antibodies for 3 hours at room temperature. Following the labeling, the sections were washed 3 times in 1x TBS buffer, 5 minutes each. In one staining (pTau and Thioflavin S), sections were incubated with Thioflavin S, followed by 2 × 5 minutes washes with distilled water. After the last wash, sections are incubated with DAPI (4′,6-diamidino-2-phenylindole, Cat. no. 10236276001 Roche, Switzerland) for 10 minutes at room temperature. Thereafter, the sections were washed 2 times, 5 minutes each with TBS, and mounted with custom-made aqueous mounting media containing 1,4-Diazabicyclo[2.2.2]octane (803456 EMD Millipore, USA). For the lipid droplet Nile red staining, sagittal brain sections were mounted on to the glass slide and air-dried, followed by a rinse with deionized water for 1-2 minutes. Nile red/glycerol staining solution (2μg/ml) was added onto the tissue, incubated for 5 minutes, and coverslipped before being examined using a confocal fluorescence microscope (Zeiss LSM 800, Germany).

### Image quantification

To quantify the glial response to an increase in systemic and central inflammatory profile, the morphology of these brain innate immune cells has been analyzed. Iba-1 positive microglia images (10 ROI/section) were taken at 40x oil immersion objective using a Confocal microscope (Zeiss LSM 800, Germany) with 1 μm Z intervals resulting in stacks of 20-30 slices at 512 × 512 pixel resolution. The soma area, perimeter, circularity, skeleton analysis was performed as instructed (Young and Morrison 2018; Davis et al. 2017) and using custom-made macros (Suppl. Materials and Methods). For the Fractal analysis on aged mice microglia 60x Oil immersion objective was used to generate in-depth microglia morphological data. Mean fluorescence intensity (pTauT231) of the CA1 field was measured in the same age group using the ImageJ ROI manager. Amyloid and Lcn2 staining were analyzed for % area coverage using the custom made ImageJ macros (Suppl. Materials and Methods). Nile red positive lipid droplets were analyzed using the Analyze particle function in ImageJ. For the histological data, ImageJ macros were created and used for the analysis for an unbiased morphological quantification (Suppl. Materials and Methods). For the western blot, protein band images were analyzed using the ImageJ Gel quantification tool.

### Statistical analysis

Homogeneity of variance for each variable within a group/ age was verified by Levene’s Test and comparative analysis performed either using the parametric test for normally distributed data (Student’s t-test) or non-parametric test for non-normally distributed data (Mann Whitney test). Statistical significance was assessed using unpaired, 2-tailed Student’s t-te*st*. Results (mean ± SEM) were obtained using Excel Real Statistics plugin. *P* values less than 0.05 were considered significant. RNA sequencing data were analyzed using the ‘R’DESEQ2 package and statistically significant transcripts were chosen if p<0.05 and p adjusted for multiple testing <0.1. For the flow cytometry data, statistical analysis was performed through unpaired Welch’s t-test using GraphPad Prism Version 8.1.2. Optical densities of immunoblots were compared between groups of the same age using Student’s t-test. Morphology measurements of microglia were analyzed using the Kolmogorov–Smirnov test and in aggregate using Student’s t-test. Correlation analysis was conducted using R and significance tested using Student’s t-test.

## Results

The goal of the study is to demonstrate that low-grade chronic inflammation, through prenatal and postnatal sterile infection with PolyI:C is sufficient to cause an AD-like pathology.

### Chronic inflammation after double PolyI:C challenge

To obtain systemic low-grade chronic inflammation we have performed double sterile infection with the TLR3 (Toll-like receptor 3) stimulant, PolyI:C, in the mother at gestational day 17 and in the offspring at 2.5 months (PP). This protocol deviates from the previous studies using either prenatal injection alone or the combination of prenatal and senior injections (Krstic et al. 2012), which could not be replicated by others (Giovanoli et al. 2015) and aims at investigating progressive pathophysiological changes in response to inflammation already in midlife. Animals were analyzed cross-sectionally at 3, 6, 9, and 16 months, taking saline-injected controls as reference (NN) (Fig. 1A). From 3 to 6 months of age levels of four circulating inflammatory cytokines (MCP-1, IL-6, TNF-α, and IL-10) are progressively higher in PP as compared to NN (Table 1). At 9 months, all 5 cytokines are increased already in the NN, as compared to the previous stages and only IL-6 is 6 fold elevated in PP mice (p<0.05) (Table 1). At 16 months, NN and PP mice show no difference in humoral immune factors (Table 1). Considering the upregulation of a large population of circulating chemokines in the blood of PP mice, we examined the neutrophils, monocytes, and total polymorphonuclear cells (PMNs) from the freshly isolated whole blood and brain samples to study any infiltration of these leukocytes into the brain. Samples were analyzed using the flow cytometry for the expression of neutrophils and monocytes using antibodies stained for Ly6G, Siglec, F4/80, CD11b. Although no changes are observed in the 3 months of age between groups, at 6 months PP mice exhibit elevated levels of neutrophils and PMNs in the systemic circulation but no significant differences are seen in the brain of PP mice as compared to NN animals (Suppl. Fig. 1A and B). On the other hand, the monocyte population differs neither in the blood nor in the brain (Suppl. Fig. 1C). While there is no trace of infiltrating immune cells in the brain at 3 or 6 months, analysis by RT-PCR of *IL-6* transcripts shows a significant increase in PP mice at 6 months, which then stabilizes at 9 months and decreases at 16 months compared to NN controls (p<0.01) (Fig. 1B). We further analyzed how the inflammatory transcripts correlate with cell-type-specific transcripts for neurons (*GRIN1,* Glutamate Ionotropic Receptor NMDA Type Subunit 1), microglia (*Iba1;* Ionized calcium-binding adaptor molecule 1) and astroglia (*GFAP;* Glial Acidic Fibrillary Protein) in PP and NN animals. We observe that *GRIN1* expression is stable except for a trending increase at 9 months (Fig. 1C), whereas *Iba1* shows dynamic changes with a progressive increase from 6 to 9 months and drop at 16 months (p<0.01) (Fig. 1D) similarly to the proinflammatory cytokines (1IL-6, r_16mo_=0.77 and IL-1*β*, r_16mo_=0.68). At 6 and 9 months also, *GFAP* expression increases but decays to normal levels at 16 months (Fig. 1E). The temporal evolution of *GFAP* overlaps with that of IL-1*β*, (r=0.62, p<0.05; Fig. 1F). Our results show that the combination of prenatal and an early postnatal PolyI:C immune activation causes a prolonged and sustained systemic inflammatory response with the recruitment of circulating leukocytes of the innate immunity, and increased neuroinflammation and gliosis at mid-age.

**Table 1:**
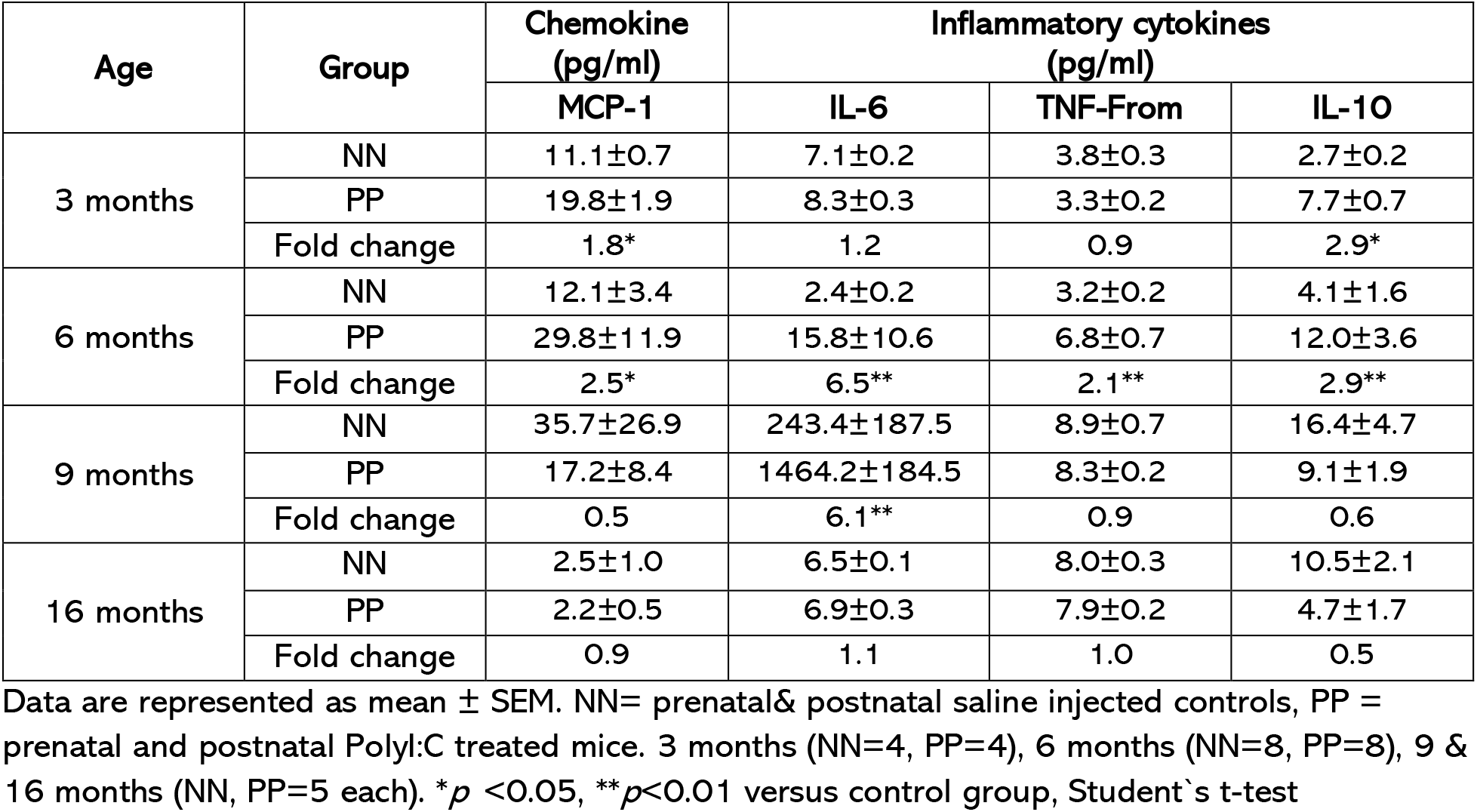
Plasma inflammatory panel in aging PP and NN mice

| Age | Group | Chemokine<br>(pg/ml) | Inflammatory cytokines<br>(pg/ml) |  |  |
| --- | --- | --- | --- | --- | --- |
|  |  | MCP-1 | IL-6 | TNF-From | IL-10 |
| 3 months | NN | 11.1±0.7 | 7.1±0.2 | 3.8±0.3 | 2.7±0.2 |
| | PP | 19.8±1.9 | 8.3±0.3 | 3.3±0.2 | 7.7±0.7 |
|  | Fold change | 1.8* | 1.2 | 0.9 | 2.9* |
| 6 months | NN | 12.1±3.4 | 2.4±0.2 | 3.2±0.2 | 4.1±1.6 |
| | PP | 29.8±11.9 | 15.8±10.6 | 6.8±0.7 | 12.0±3.6 |
|  | Fold change | 2.5* | 6.5** | 2.1** | 2.9** |
| 9 months | NN | 35.7±26.9 | 243.4±187.5 | 8.9±0.7 | 16.4±4.7 |
| | PP | 17.2±8.4 | 1464.2±184.5 | 8.3±0.2 | 9.1±1.9 |
|  | Fold change | 0.5 | 6.1** | 0.9 | 0.6 |
| 16 months | NN | 2.5±1.0 | 6.5±0.1 | 8.0±0.3 | 10.5±2.1 |
| | PP | 2.2±0.5 | 6.9±0.3 | 7.9±0.2 | 4.7±1.7 |
|  | Fold change | 0.9 | 1.1 | 1.0 | 0.5 |

Data are represented as mean ± SEM. NN= prenatal& postnatal saline injected controls, PP = prenatal and postnatal PolyI:C treated mice. 3 months (NN=4, PP=4), 6 months (NN=8, PP=8), 9 & 16 months (NN, PP=5 each). \* $p$ <0.05, \*\* $p$ <0.01 versus control group, Student's t-test

### Progressive tauopathy in PP mice

Inflammatory mechanisms are central to AD pathology involving both the interaction between inflammatory stimuli, proinflammatory cytokines/mediators, and AD pathological hallmarks such as aggregates of amyloid-beta (Aβ) and hyperphosphorylated tau protein (p-tau) (Kinney et al. 2018), underlying neural network dysfunction and brain atrophy. We determined the impact of systemic immune challenge on the pre-tangle pathology by analyzing tau hyperphosphorylation at positions 231 and 205 using western blot (Fig. 2A) and immunohistochemistry (Fig. 2F and Suppl. Fig. 2A) on hippocampal tissue from 3 to 16 months of age. From the immunoblot, the content of hippocampal p-tau (pTau T231) relative to total tau (ptau/tau) increases dramatically in PP mice at 6 months, remains elevated at 9 months and then is indistinguishable from NN mice at 16 months (Fig. 2B and 2C). The progressive tauopathy is confirmed by immunolabeling of pTau fibers (pTau T205) in the hippocampal CA1 region and CA1 stratum lacunosum (Suppl. Fig. 2A), with a significant increase in pTau pixels in the CA1 region at 16 months (Suppl. Fig. 2B). Tau hyperphosphorylation and the formation of *β*-sheet fibrils, labeled by Thioflavin-S, can be observed in the CA3 field at 9 months in PP mice, in contrast to NN (Fig. 2F). To assess whether the progressive tauopathy in PP is associated with synaptic abnormalities and neuroinflammatory responses, we quantified the levels of synaptophysin and GFAP overtime (Fig. 2D). At 16 months in PP mice, we observe a drop in synaptophysin expression accompanied by an increase in GFAP (Fig. 2E). The increase in GFAP immunoreactivity is confirmed by immunohistochemistry of the CA3 region showing the presence of amyloid-*β*_1-42_ labeled by an antibody recognizing the amino acid region 1-16 (A**β**1-16) and p-tau aggregates (Fig. 2G). Further analysis of amyloid-*β* fibrils in the 16 months old mice indicates that A*β*_1-42_ is also present in NN old mice with internalization in GFAP positive glial cells, which is more pronounced in PP mice (Fig. 3A, white stars). In addition in 16 months PP mice, A*β*_1-42_ positive aggregates are visible in vessels ensheathed by astroglial endfeet reflecting a cerebral amyloid angiopathy (CAA) (Fig. 3A, white arrows). Increased colocalization between insoluble A*β*_1-42_ and p-tau in 16 months old PP as compared to NN both in the hippocampus (Fig. 3A) and the entorhinal cortex (Fig. 3B). In the latter region, A*β*_1-42_/p-tau positive aggregates display a stellate morphology resembling core-plaques (Fig. 3C, insert). The number of plaques rises 4 folds in PP as compared to NN (Fig. 3D), with a 70% larger amyloid-*β* burden in the area examined (Fig. 3E). Quantification of soluble A*β*1-42 from the entorhinal cortex shows a peak in A*β*_1-42_ release at 6 months in PP mice which later subdues (Fig. 3F), possibly as a result of insoluble aggregate formation in the brain.

**Figure 2.**
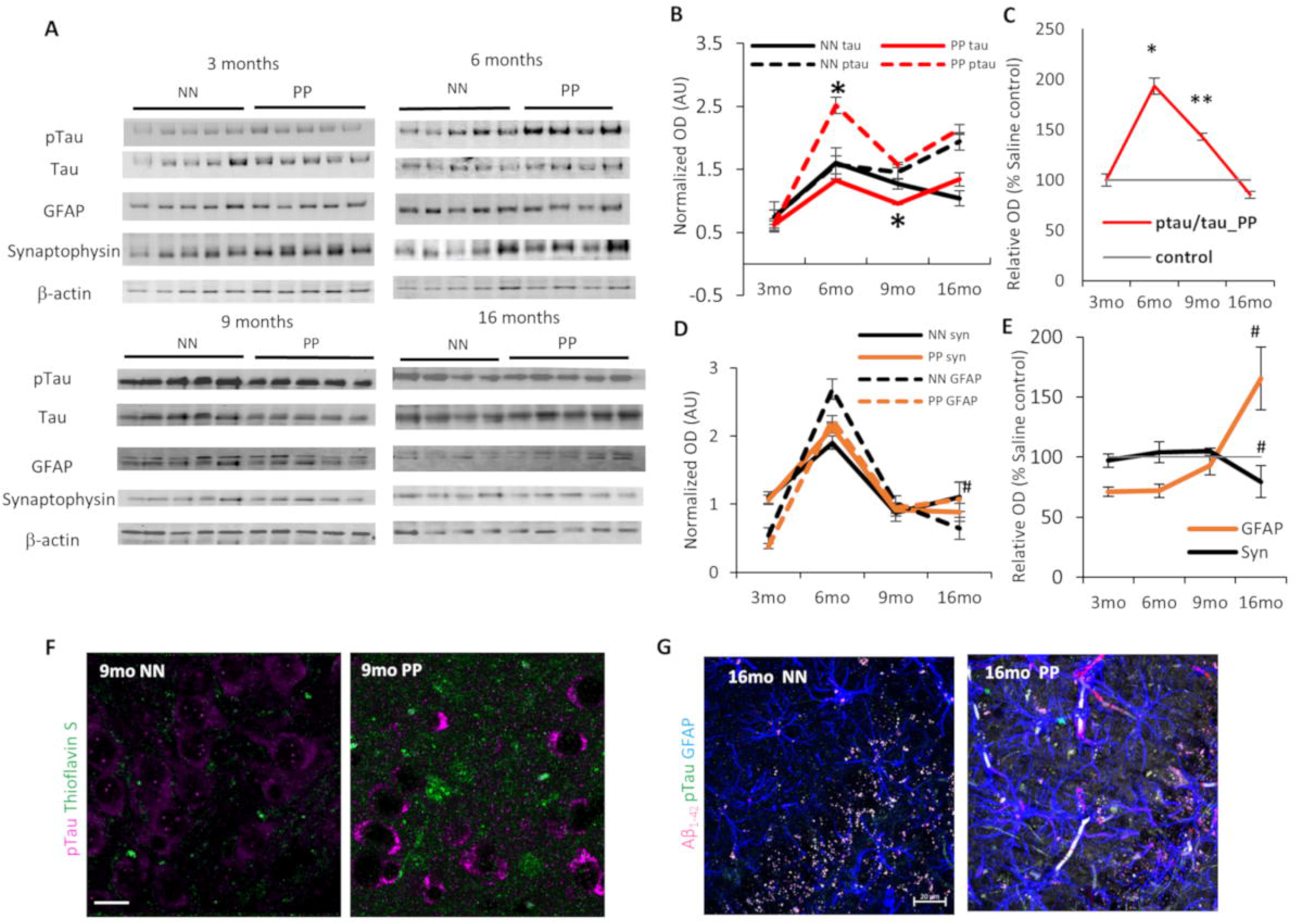
The Effect of PolyI:C on p-tau accumulation in the hippocampus. **(A)** Immunoblots of hippocampal lysate from PP and NN mice at 3, 6, 9, and 16 months, for p-tau, tau, GFAP, Synaptophysin. β-Actin was used as housekeeping control. **(B, D)** Relative quantification of ptau, tau, GFAP, and Synaptophysin represented in normalized optical density. **(C, E)** Percentage of relative changes of these proteins compared to the saline treated controls (NN). Representative immunostaining for **(F)** p-tau, Thioflavin S (9 months), and **(G)** Aβ_1-42_, p-tau, and GFAP at 16 months age group. Scale bar in H is 45 μm, J is 20μm. Values represented as mean ± SEM, n=4-5 mice per age, and treatment. *p<0.05, **p<0.005, #p=0.07. Student’s t-test

**Figure 3.**
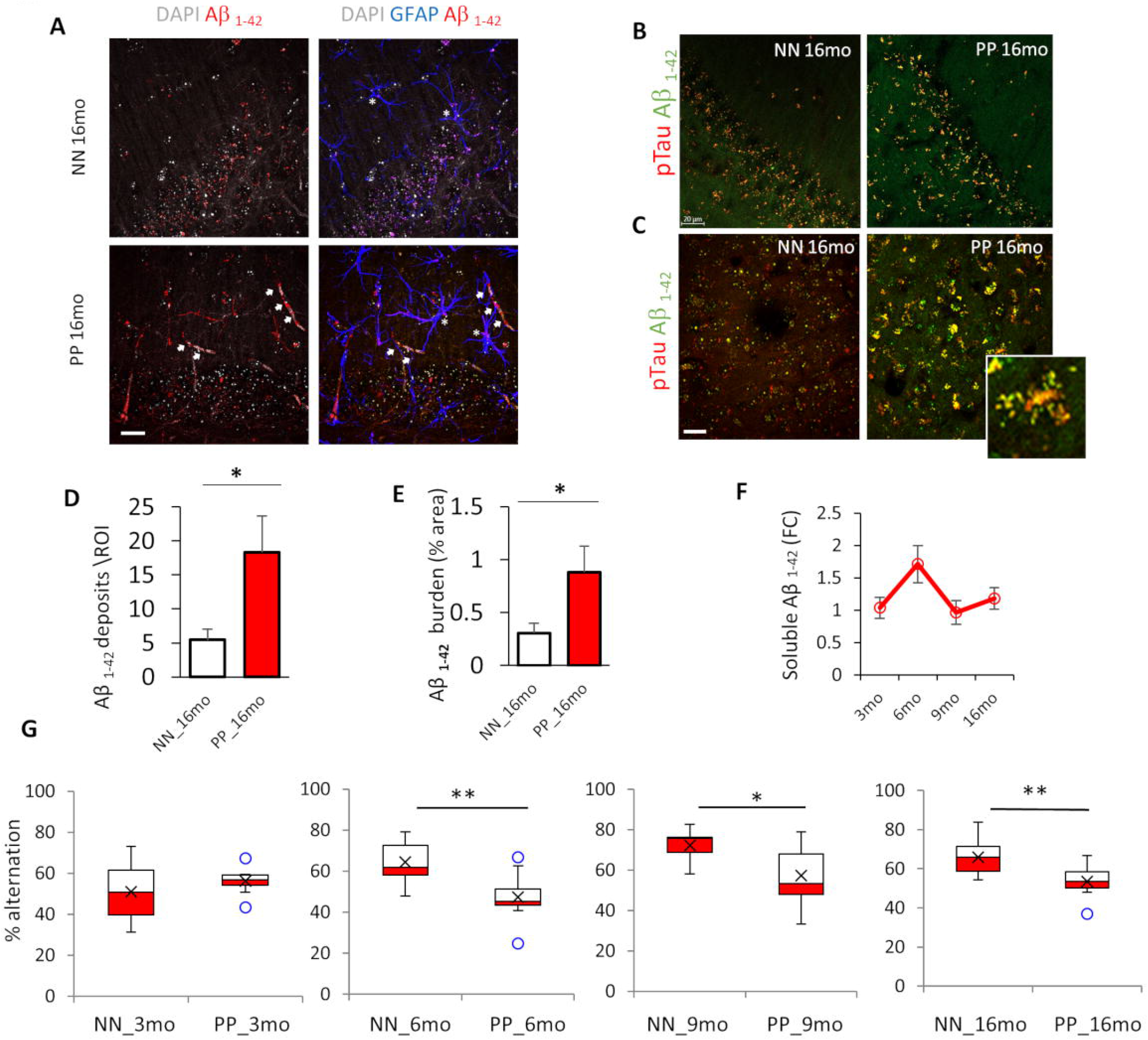
Amyloid fibrils in the hippocampus and entorhinal cortex with spatial memory deficits. **(A**) Double immunolabeling for Aβ_**1-42**_ and GFAP shows internalization of amyloid in GFAP+ glia cells (stars) in NN and PP mice at 16 months and strong depositions of amyloid in vessels (white arrows) in PP mice. **(B**) Representative immunostaining of Aβ_**1-42**_ and pTau^231^ in the hippocampus and **(C**) Entorhinal cortex of aged 16 months PP & NN controls. Insert showing p-Tau & Aβ_**1-42**_ colocalization. (**D**) Quantification of amyloid aggregates >5μm^2^ and (**E**) percentage area covered per region of interest (n=5-6 mice per condition & 4-5 ROI per mice). (**F**) Immunobead assay showing dynamic trend in soluble Aβ_42_ level fold changes in the cortical lysates relative to control (n=4-6 mice per age group & condition). **(G)** Box plots indicate a significant impairment in the percent of spontaneous alternation entry in PP mice compared from 6 to 16 months as compared to NN (n=7-10 mice per treatment and age). Values in bar charts and line graph are represented as mean ± SEM. *p<0.05, **p<0.005 Student’s t-test. Scale bar in A and B=20μm.

### PolyI:C-induced memory impairment

Based on the proteinopathy in the hippocampal and entorhinal cortex region following double PolyI:C injection, we assessed whether spatial working memory is affected. Previous work has shown that spontaneous alternation, a form of working memory, is entirely dependent on hippocampal synaptic function (Pioli et al. 2014; McHugh et al. 2008). PP and NN animals were subjected to the Y-maze spontaneous alternation task at 3, 6, 9, and 16 months. The number of arm entries and sequence were recorded, and the percent alternation was calculated (Fig. 3G). Although the activity of PP treated animals increases in terms of alterations at 3 months, the percentage of alternation did not change at this age as compared to NN (t17 = 0.76, P = 0.46 n = 10 for Control and n=9 for PolyI:C). On the other hand, deficits in the spontaneous alternation are observed in the PP group both *at* 6 months (t17 = 3.21, p = 0.005, n = 10 for NN and n=9 for PP), 9 months (t15 = 2.57, p = 0.013, n = 9 for NN and n=8 for PP) and 16 months (t13 = 2.57, p = 0.02 n = 8 for NN and n=7 for PP). The number of arm entries in the Y maze task is not significantly different across all the age groups (Suppl. Table 4) indicating normal locomotory behaviour representing a non-confounding factor for the percentage of an alternation. However, from 9 months of age both PP and NN display about half of the number of entries as compared to younger animals. Since early and late gestational PolyI:C treatment has been linked to stress and anxiety-like phenotypes relevant to schizophrenia (Hui et al. 2018; Silveira et al. 2017) we tested these mice for Light/dark, Elevated O-maze at 3 and 6 months and Open field tasks at all ages. We found no difference in the anxiety-like behaviour (Suppl. Table 5) disambiguating the spatial memory impairments observed. Together the data suggest that prenatal and early postnatal viral-like immune activation through PolyI:C treatment causes proteinopathy of the limbic regions with sustained working memory impairment.

### Activation and phenotypic change in microglia of PP animals

Earlier studies have shown that systemic infections and the subsequent peripheral immune activation have a strong effect on brain function, glial response representing a risk for dementia (Cunningham and Hennessy 2015; Cunningham 2013). Microglia cells are the primary phagocytic innate immune cells of the brain that get stimulated upon immune activation. They normally exist in a resting state displaying a ramified morphology, while in intermediate states display a bipolar or rod-like phenotype (Davis et al. 2017) and in an activated state an amoeboid, irregular shape (Ling and Wong 1993). Quantification of Iba-1 positive cells per mm^3^ in the immunostained hippocampus (Fig 4A), indicates no difference between PP and NN at the different stages (Suppl. Table 6). Next, we assessed whether the PP brains show altered microglial cell morphologies reflecting an activated inflammatory status. We quantified morphological parameters like soma size, perimeter, circularity, skeleton analysis of Iba-1 positive microglia (Fig. 4), and finally we conducted the fractal analysis in NN & PP mice across staging. From 6 months of age, Iba-1 positive microglia show an increased ramified morphology as detected by immunofluorescence microscopy (Fig. 4A). Quantitative soma analysis shows a slight increase in the soma size, typical of activated microglia in PP mice at 3 months (22%, p<0.005), which last up to 9 months (58%, p<0.005 (Fig. 4B). The cell soma perimeter measurement shows dynamic changes at 3 months (11% increase, p<0.01), and a significant increase at 9 months (57%, p<0.005), while no changes are detected at 6 and 16 months (Fig. 4B). Circularity, which indicates the roundness index, shows a deviation from circularity in the intermediate ages at 6 months (10% decrease, p<0.005), 9 months (21% decrease, p<0.005) and 16 months (10% increase, p<0.05), indicating signs of progressive microglial activation with aging (Fig. 4B). These dynamic changes were further analyzed using the distribution curve analysis which indicates an overall shift in the number of cells with an increase in microglial cells soma size and perimeter (Suppl. Fig. 3A-3B) with irregularly shaped cell body during the adult and mid-aged intermediate stages (6 and 9 months) while at later stages (16 months), microglia become more circular or rounded suggesting morphological fluctuations across staging and signs of microglial activation (Suppl. Fig. 3C). To further characterize the phenotypic changes in microglia, reflecting their activation state, we performed the skeletal analysis of Iba-1 positive microglia, at 3, 6,9, and 16 months in PP and NN mice. Skeletonized Iba-1 renderings reveal a significant increase in the number of endpoints/cell, maximum branch length/cell, and branch length/cell (Fig. 4C) in PP mice with age groups 3 months to 9 months indicating hyper ramified microglia. However, at 16 months, microglia in PP brains show a significant drop in the number of endpoints/cells and an overall decrease in the branch and maximum branch length (Fig. 4C). Therefore, prenatal and early postnatal systemic PolyI:C immune activation induces hyper ramified microglia in the offspring which later undergoes a transition from hyperactivated to more bushy or amoeboid with the reduction in microglial endpoints.

**Figure 4:**
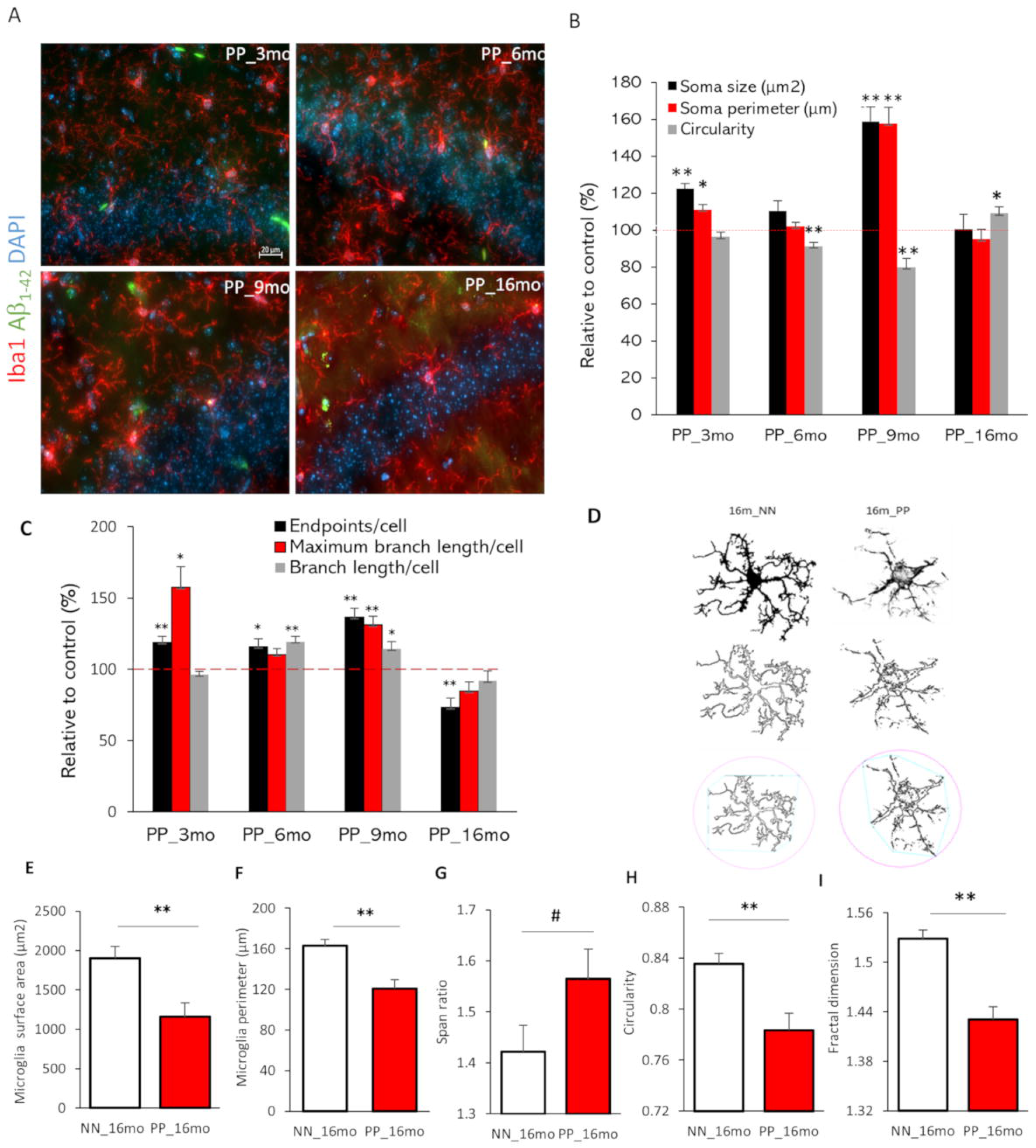
Longitudinal changes in hippocampal microglia. **(A)** Immunofluorescence staining for Iba1 and Aβ staining in the PolyI:C treated mice. **(B)** Dynamic changes in microglia soma size, perimeter and circularity were quantified in PolyI:C (PP) mice in different age groups. Data represented as % relative to controls. (n= 150-180 cells from 3-4 mice per group & condition in 3m, 6m stages; n=85 cells in 9m and n=55 cells at 16 months from 3 mice). **(C)** Bar chart of % relative values relative to controls for the average endpoints, maximum branch length, and branch length per cell. Red dotted line indicates the 100 percent control line (n=55-65 ROIs/3, 6, 9 months group and 25 ROIs/16 months; n=3 mice per age group and condition). **(D)** Example of fractal analysis of microglia from the hippocampus of PolyI:C and saline treated 16 months aged mice. Single-cell fractal analysis indicates the surface area **(E)** and perimeter **(F)** is decreased in PP mice, in contrast, the span ratio **(G)** showed a higher trend. PP mice showed a significant decrease in circularity **(H)** and fractal dimension **(I)** (n=45 cells from 3 mice per group and condition). Values in the bar charts are represented as mean ± SEM. #p=0.07,*p<0.05, **p<0.005 Student’s t-test. Scale bar, A= 20μm.

Having detected a microglia morphological transition in aged PP mice, confocal imaged microglia cells were further investigated using fractal analysis (FracLac plugin of ImageJ) which renders the shape of microglia cells and quantifies the parameters of cell area, cell perimeter, Span ratio, circularity, and fractal dimension, the latter reflecting pattern complexity. The fractal analysis revealed significant changes in hippocampal microglia parameters in the aged 16 months PP mice versus NN (Fig. 4D). Normal resting microglia are complex with a higher fractal dimension. In the PP mice total microglia cell surface area, the perimeter is reduced indicating a more compact shape (Fig. 4E-4F). Fractal dimension as a measure for complexity and circularity as a measure for the roundness is also reduced after immune activation (Fig. 4H-4I). On the other hand, the span ratio is increased reflecting microglia elongation (ratio of cell length and width) (Fig. 4G). Span ratio and circularity are inversely proportional indicating an activated microglia state. Altogether, systemic inflammation through pre- and post-natal PolyI:C induces profound changes in microglia morphology indicating a shift from resting to an activated state in aged PP animals.

### Dynamic remodeling of the hippocampal transcriptome

To understand the mechanisms underlying the spatial memory deficits and the pathophysiological processes associated with the proteinopathy, and neuroinflammation we performed a cross-sectional hippocampal bulk mRNA sequencing on 3, 6, 9, and 16 months PP and NN animals. The analysis reveals many differentially expressed genes (DEGs) in the hippocampus (log2 fold change cut-off:05, padjusted <0.05) between treatments. The gene expression profiles as visualized in the volcano plot (Fig. 5A-5D), show that in aged animals differential expression is highest with a peak at 9 months, coinciding with the neuroinflammatory switch in microglia cells. A GO analysis contextualized to the synapse (SYNGO) (Koopmans et al. 2019) indicates a dynamic shift in DEGs from the presynaptic compartment at 3 months (Supp. Fig 4A) to the postsynaptic terminal at 9 and 16 months (Supp. Figure 4C and 4D), with a non-synaptic stage at 6 months (Supp. Fig 4B). At 3 months, out of 35 DEGs between PP and NN, 21 are downregulated, and 14 are upregulated (Fig. 5A). Gene ontology enrichment analysis (GEA) of the biological processes using a 5% false discovery rate indicates general repression in genes associated with synaptic transmission, calcium signaling, extracellular matrix organization, and secretion (Suppl. Table 7). Pathway analysis based on a composite KEGG, Reactome, WikiPathways dataset shows high interconnectivity between cellular cascades associated with extracellular matrix organization, chemical synaptic transmission, and calcium signaling (Suppl. Fig 5A). In the 6 months, out 32 DEGs between PP and NN (Fig. 5B), 22 are downregulated, and 10 upregulated. A GEA indicates ongoing processes of morphogenesis and gliogenesis (Suppl. Table 7), with highly overlapping pathways (Suppl. Fig. 5B). At 9 months (Fig. 5C) out of 196 DEGs, 90 are downregulated, and 106 are upregulated. The GEA shows enrichment in several processes associated with vascular remodeling, neurogenesis, morphogenesis, cytokine response, and cell death (Suppl. Table 7). Pathway analysis indicates partially overlapping cellular cascades shared among focal adhesion, BMP signaling, MAPK signaling, and neuronal injury (Suppl. Fig. 5D) confirming the concurrent processes of morphogenesis and cell death at this stage. In the 16 months PP mice (Fig. 5D) from the 99 DEGs, 50 genes are downregulated and 49 genes are upregulated. GEA of the DEGs indicates enrichment in the response to metal ions, the reactive oxygen response, morphogenesis, the regulation of cellular proliferation, and the response to hormones (Suppl. Table 7). Pathways analysis shows less interconnected cascades implicated in potassium ion transmembrane transport, monocytes proliferation, lamellipodium organization, and glucose metabolism (Suppl.Fig. 5D).

**Figure 5:**
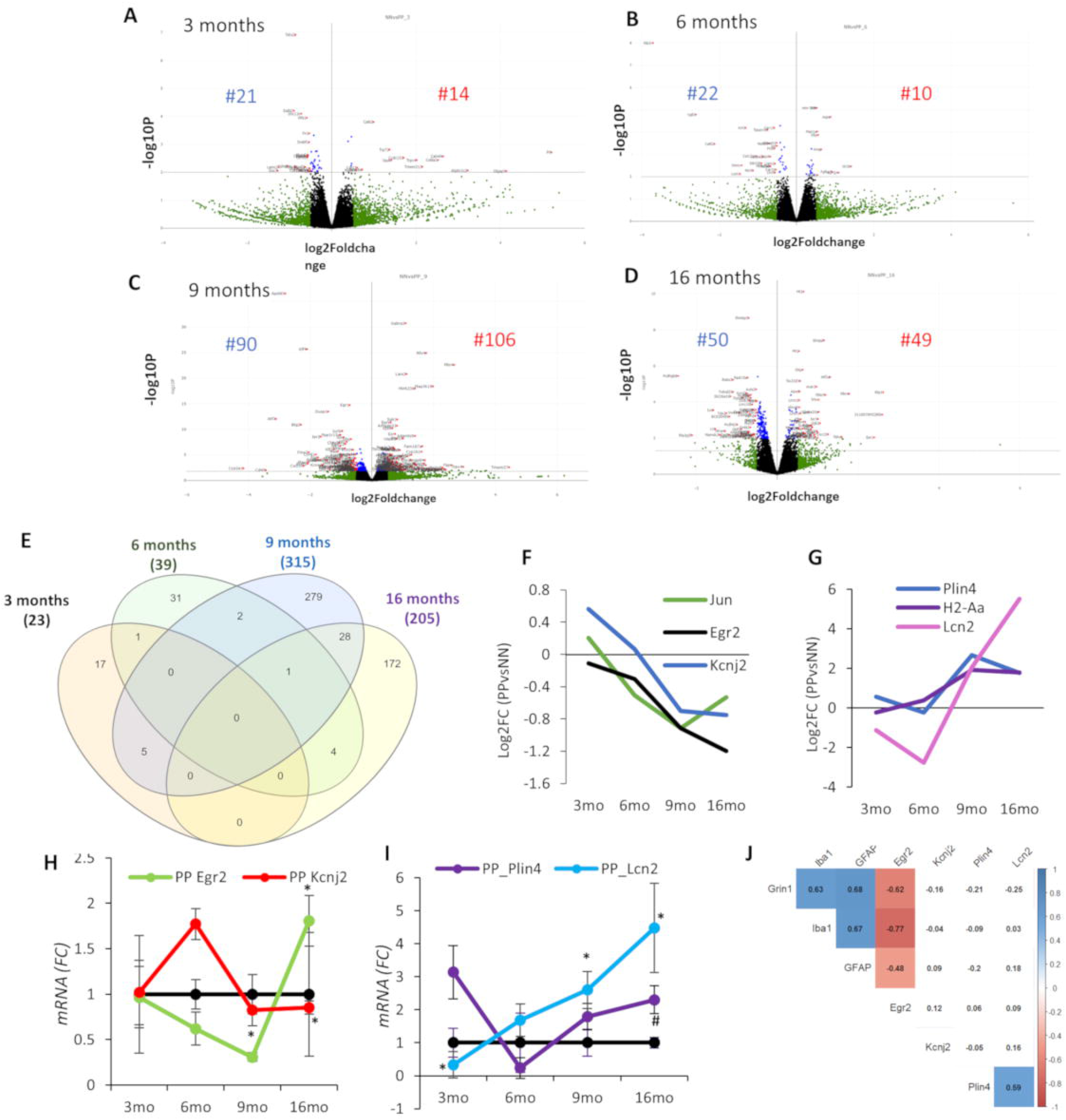
Bulk transcriptomics analysis of the hippocampus in the aging PolyI:C. Gene expression changes after the prenatal and early postnatal immune challenge. Volcano plot with log2-fold change (X-axis) and −log10 p-value (Y-axis) at different age groups **(A)** 3 months **(B)** 6months **(C)** 9 months and **(D)** 16 months in PP versus NN controls. Genes with log2 fold changes >0.5 are shown in green. Significantly regulated genes are shown in red (log2 fold change >0.5, p<0.05, left downregulated, and right upregulated). Genes with insignificant log2 fold changes <0.5 are shown in black [controls (NN) n=3/age, PolyI:C (PP) n=3/age]. **(E)** Venn diagram indicating the number of uniquely and commonly affected genes in the aging hippocampus following prenatal and early postnatal PolyI:C (PP) treatment. Few common genes along with their Log2 Fold changes either **(F)** downregulated or **(G)** upregulated were plotted across staging. Transcript validation analysis for **(H)** *Egr2* and *Kcnj2* and **(I)** *Plin4* and *Lcn2*. **(J)** Correlation matrix with highlighted significant association from 9 to 16 months among the differentially expressed cell-type markers (*GFAP, Iba1,* and *Grin1*) and *Egr2*, *Kcnj2*, *Plin4* and *Lcn2* between PolyI:C and saline controls [controls (NN), n=5/age, PolyI:C (PP) n=4-5/age]. Line plots show values as mean ± SEM. *=p<0.05, **=p<0.01, #=p@0.1, Student’s t-test.

To better understand the dynamic changes in the sterile infection model undergoing with age, we compared the significantly differentiated genes between PP and NN at the cross-sectional time points. As shown in the Venn diagram (Fig. 5E), the number of common genes that are uniquely and commonly affected in the hippocampus of 3, 6, 9, 16 months PP mice indicates a large number of overlapping gene sets between 9 and 16 months, while fewer genes are shared with earlier stages. In particular, 6 shared genes can be subdivided into 2 categories reflecting a progressive cell-communication dysfunction (I) and a proinflammatory drive (II) in aging PP mice. (I) Genes downstream of MAPK-signalling (*c-Jun*) (E. K. Kim and Choi 2010), NFKB-signaling (*Egr2;* Early growth response protein 2) (Williams et al. 1995), responsible for neuronal excitability (*Kcnj2;* Potassium Inwardly Rectifying Channel Subfamily J Member 2) (Binda et al. 2018), with reported association with synaptic dysfunction, are downregulated in PP mice starting from 6 months (Fig. 5F). (II) On the other hand, genes related to neuroinflammation such as the lipid-droplet dependent gene (*Plin4;* Perilipin 4) (Han et al. 2018), pro-inflammatory genes (*H2-Aa*; Immunohistocompatibility-complex) (Van Hove et al. 2019) and acute-phase proteins regulating to the inflammatory response (*Lcn2*; Lipocalin-2) (Dekens et al. 2020) are upregulated in aging PP mice, starting from 9 months (Fig. 5G). Overall, the gene remodeling across the aging continuum replicates processes typical of AD as neuronal network breakdown, altered immune response, chronic neuroinflammation and vascular dysfunction.

### Substantial changes in brain metabolism and inflammation

We further validated the RNA-seq analysis via RT-PCR, on some of the significantly DEG with a reported association to AD as well as genes belonging to the neuronal (I) (Fig 5H) and metabolic (Fig 5I) gene groups. At 3 months, *Cacna1g* (Calcium Voltage-Gated Channel Subunit Alpha1 G), which was previously reported to decay with aging and regulate amyloid-*β* production (Rice et al. 2014), is unchanged (0.9) in contrast to the observed reduction via RNA-seq (Log2FC=−0.55), suggesting that aggregate changes rather than single-gene changes may contribute to the modeled conductivity dysfunction (Suppl. Tab. 6). As expected at this stage, *Plin4* and *Egr2* are unchanged, matching the RNAseq data (Table 2). At 6 months, we analyzed one of the genes with the strongest downregulation at the RNA-seq, with reported association with cognitive impairment in chronic cerebral hypoperfusion (Xie et al. 2018). The *Glpr2* (Glucagon-Like Peptide 2 Receptor) decrease (80%), is confirmed (Table 2; p<0.05). Also, *Ide* (Insulin-degrading enzyme), which has been implicated in the clearance of insulin and amyloid-β (Qiu and Folstein 2006), show a comparable increase, between RNA-seq and RT-PCR analysis, but did not reach significance in the RT-PCR result (Table 2; p=0.16). Interestingly, *Kcnj2* show a significant 2 fold increase at 6 months (Table 2; p<0.05), suggesting a modulatory K+ currents effect. *Plin4* remain unchanged between PP and NN at 6 months in both analyses (Table 2). At 9 months, the reduction in *c-fos*, *c-Jun*, *Notch1*, *Kcnj2*, *Egr2*, and the increase in *Lcn2* can be confirmed at the RT-PCR, while *Plin4* show a non-significant increase using RT-PCR (Table 2; p=0.24). At 16 months, *Lcn2, Plin4* trends are reproduced according to the RT-PCR, while *Kcnj2* shows a comparable but not significant decrease (Table 2; p=0.36). The data indicates that around ¾ of the RNA-seq outputs could be replicated via RT-PCR validation, confirming the robustness of the bulk RNAseq discovery method and emphasizing that aggregate dataset can explain ongoing cellular and molecular processes in such models. Validated DEG profiles of *Kcnj2*, *Egr2,* and *Plin4* and *Lcn2* are matched with the expression of genes specific for neurons (*Grin1*), microglia (*Iba1*) and astroglia (*GFAP*) at 9 and 16 months to investigate associations of the selected genes in specific cell types (Fig. 5J). *Grin1*, *Iba1,* and *GFAP* are positively associated in their peaking trend at these time points (Fig. 1C-E) supporting the neuro-glia interplay. On the other hand, *Egr2* which decreases at 9 months and increases at 16 months is inversely correlated to *Grin1*, *Iba1,* and *GFAP* suggesting a potential ubiquitous regulation of this transcription factor on the cell-type-specific changes. Finally, *Plin4* and *Lcn2* are positively associated with their increasing trend from 9 to 16 months, confirming the RNAseq data and supporting the subsequent investigation of these neuroinflammatory markers in aged PP mice. Based on downregulation of *Glpr2* in adulthood, indicating vascular hyperperfusion (Xie et al. 2018) and the subsequent upregulation of the vascular and inflammatory markers, *Lcn-2*, at 9 months, we investigated vascular integrity by validating the gene expression of Kruppel-like factor 4, *Klf-4*, at 9 and 16 months and observed a transient downregulation at 9 months (Table 2). At the same time point, angiogenesis markers such as Cytochrome P450 Family 1 Subfamily B Member 1, *Cyp1b1,* and Angiopoietin-like 4, Angptl4, with reported function in vascular AD and homeostasis (Chakraborty et al. 2018; Ghosh et al. 2016) showed a peak at 9 months, supporting pathological vascular processes in aged adult PP mice.

**Table 2.**
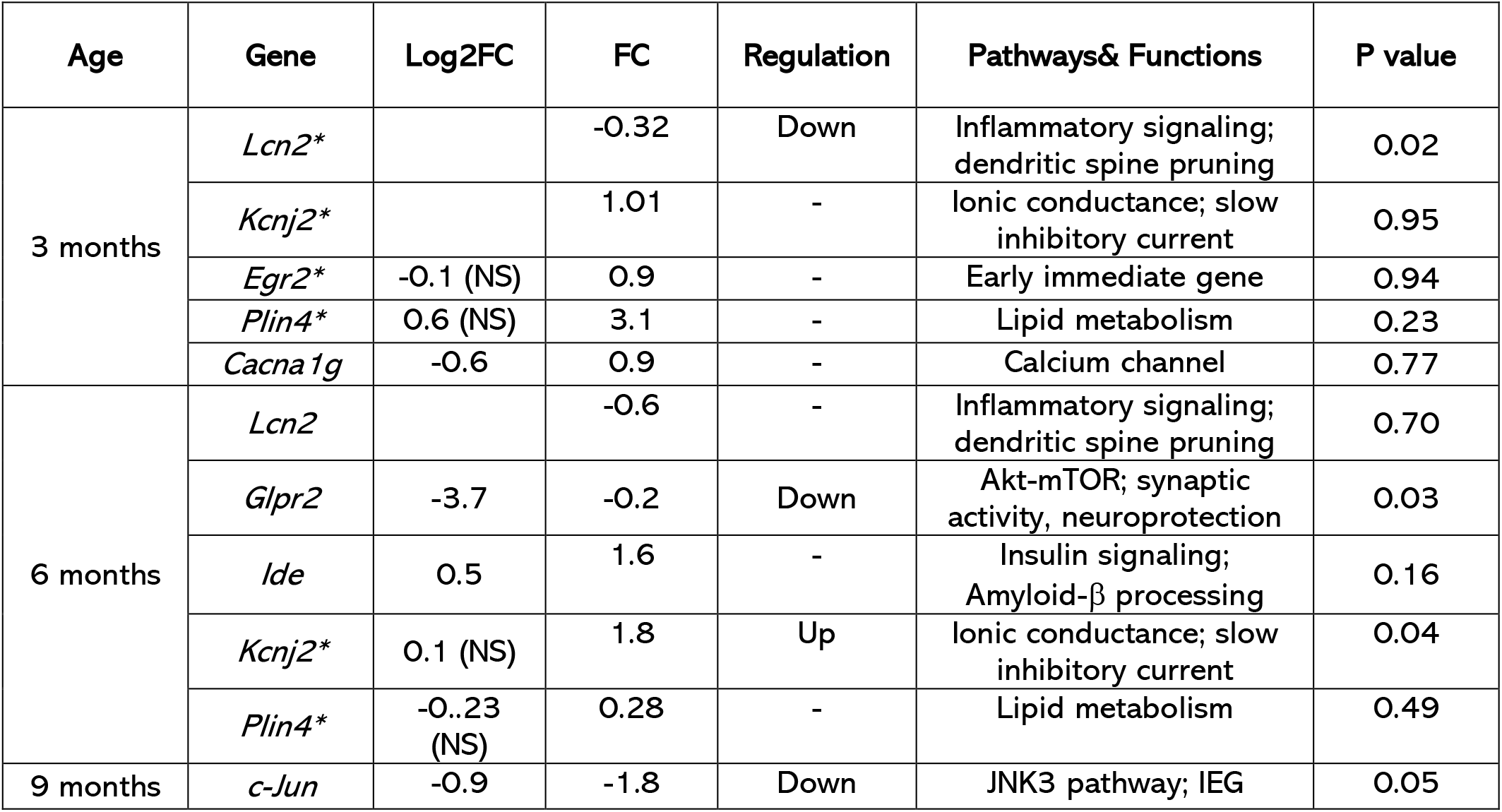
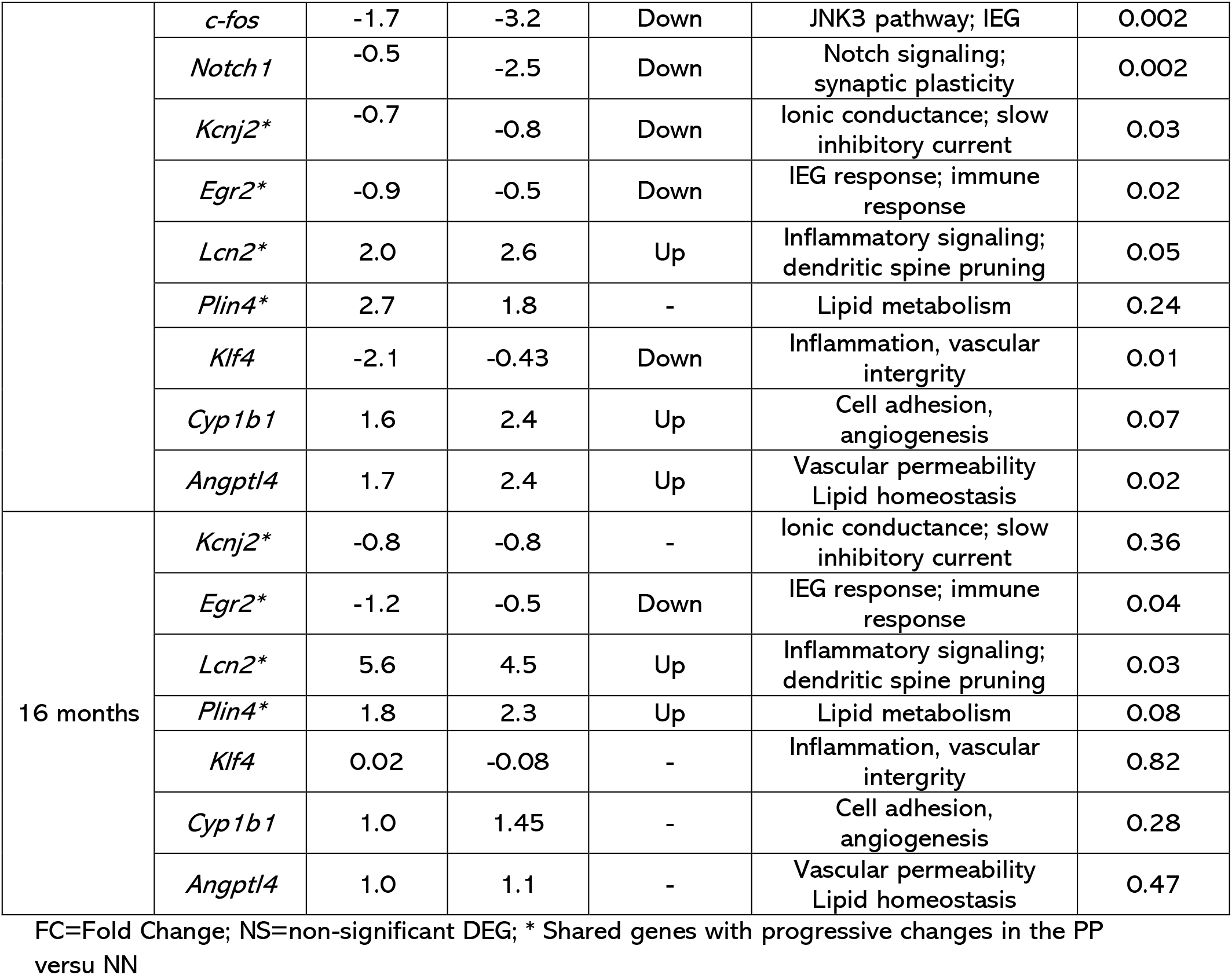
Validation of transcripts via qPCR in aging PP mice as compared to age-matched NN.

| Age | Gene | Log2FC | FC | Regulation | Pathways& Functions | P value |
| --- | --- | --- | --- | --- | --- | --- |
| 3 months | *Lcn2* * |  | -0.32 | Down | Inflammatory signaling; dendritic spine pruning | 0.02 |
|  | *Kcnj2* * |  | 1.01 | - | Ionic conductance; slow inhibitory current | 0.95 |
|  | *Egr2* * | -0.1 (NS) | 0.9 | - | Early immediate gene | 0.94 |
|  | *Plin4* * | 0.6 (NS) | 3.1 | - | Lipid metabolism | 0.23 |
|  | *Cacna1g* | -0.6 | 0.9 | - | Calcium channel | 0.77 |
| 6 months | *Lcn2* |  | -0.6 | - | Inflammatory signaling; dendritic spine pruning | 0.70 |
|  | *Glpr2* | -3.7 | -0.2 | Down | Akt-mTOR; synaptic activity, neuroprotection | 0.03 |
| | *Ide* | 0.5 | 1.6 | - | Insulin signaling; Amyloid-β processing | 0.16 |
|  | *Kcnj2* * | 0.1 (NS) | 1.8 | Up | Ionic conductance; slow inhibitory current | 0.04 |
|  | *Plin4* * | -0.23 (NS) | 0.28 | - | Lipid metabolism | 0.49 |
| 9 months | *c-Jun* | -0.9 | -1.8 | Down | JNK3 pathway; IEG | 0.05 |
|  | *c-fos* | -1.7 | -3.2 | Down | JNK3 pathway; IEG | 0.002 |
|  | *Notch1* | -0.5 | -2.5 | Down | Notch signaling;<br>synaptic plasticity | 0.002 |
|  | *Kcnj2\** | -0.7 | -0.8 | Down | Ionic conductance; slow<br>inhibitory current | 0.03 |
|  | *Egr2\** | -0.9 | -0.5 | Down | IEG response; immune<br>response | 0.02 |
|  | *Lcn2\** | 2.0 | 2.6 | Up | Inflammatory signaling;<br>dendritic spine pruning | 0.05 |
|  | *Plin4\** | 2.7 | 1.8 | - | Lipid metabolism | 0.24 |
|  | *Klf4* | -2.1 | -0.43 | Down | Inflammation, vascular<br>intergrity | 0.01 |
|  | *Cyp1b1* | 1.6 | 2.4 | Up | Cell adhesion,<br>angiogenesis | 0.07 |
|  | *Angptl4* | 1.7 | 2.4 | Up | Vascular permeability<br>Lipid homeostasis | 0.02 |
| 16 months | *Kcnj2\** | -0.8 | -0.8 | - | Ionic conductance; slow<br>inhibitory current | 0.36 |
|  | *Egr2\** | -1.2 | -0.5 | Down | IEG response; immune<br>response | 0.04 |
|  | *Lcn2\** | 5.6 | 4.5 | Up | Inflammatory signaling;<br>dendritic spine pruning | 0.03 |
|  | *Plin4\** | 1.8 | 2.3 | Up | Lipid metabolism | 0.08 |
|  | *Klf4* | 0.02 | -0.08 | - | Inflammation, vascular<br>intergrity | 0.82 |
|  | *Cyp1b1* | 1.0 | 1.45 | - | Cell adhesion,<br>angiogenesis | 0.28 |
|  | *Angptl4* | 1.0 | 1.1 | - | Vascular permeability<br>Lipid homeostasis | 0.47 |

FC=Fold Change; NS=non-significant DEG; \* Shared genes with progressive changes in the PP versu NN

Nevertheless, an increase in Lcn2 has been reported in both AD brains (Dekens et al. 2018; Naudé et al. 2012) and CSF from vascular dementia patients (Cerebrospinal Fluid) (Llorens et al. 2020a) raising the possibility that this molecule can capture a mixed vascular-AD pathology. In line with our PolyI:C model of sterile infection, the increase in Lcn-2 is likely attributed by the production and release by activated microglia, reactive astrocytes, neurons, and endothelial cells in response to inflammatory and infectious insults (Jha et al. 2015). Our immunofluorescence analysis using an antibody specific for Lcn2 shows an increase in Lcn2 protein expression in the hippocampal CA3 field of the PolyI:C mice at 9 months and 16 months aged mice (Fig. 6A). At 16 months, small A*β*_1-42_ aggregates are visible in the PP hippocampus in close association with Lcn2 positive cells (Fig, 6A, insert). Quantitative analysis of the Lcn2 signal shows a significant increase in the % of the Lcn2 stained area in PP mice in both age groups (Fig. 6B). To study lipid metabolism and intracellular lipid droplets accumulation, as a sign of neuroinflammation with aging, we have utilized the dye Nile Red which accumulates in lipids and emits red fluorescence (Greenspan, Mayer, and Fowler 1985). We observed more Nile Red-positive lipid droplets (LDs) in the hippocampal CA3 field with aging, which is even more evident in PP mice as compared to saline controls (Fig. 6C). Triple labeling with Nile Red, Neurofilament L-200, and Iba1, shows that both NF200 positive neurons and Iba1 positive microglial cells have increased lipid droplets (Fig 6D). Quantitative analysis of the LDs, represented as fold change between the PP and NN at the different time points indicates that the density of the LDs is significantly greater in PP mice starting from 6 months, resulting in increased stained area, peaking at 6 months (Fig. 6E). On the other hand, LDs’ size shows a small but significant expansion (30%) in PP mice at 3months but afterward remains unchanged between conditions (Fig. 6E). The histo-anatomical analysis confirms the presence of neuroinflammatory markers that contribute to an AD-like neuropathological progression.

**Figure 6:**
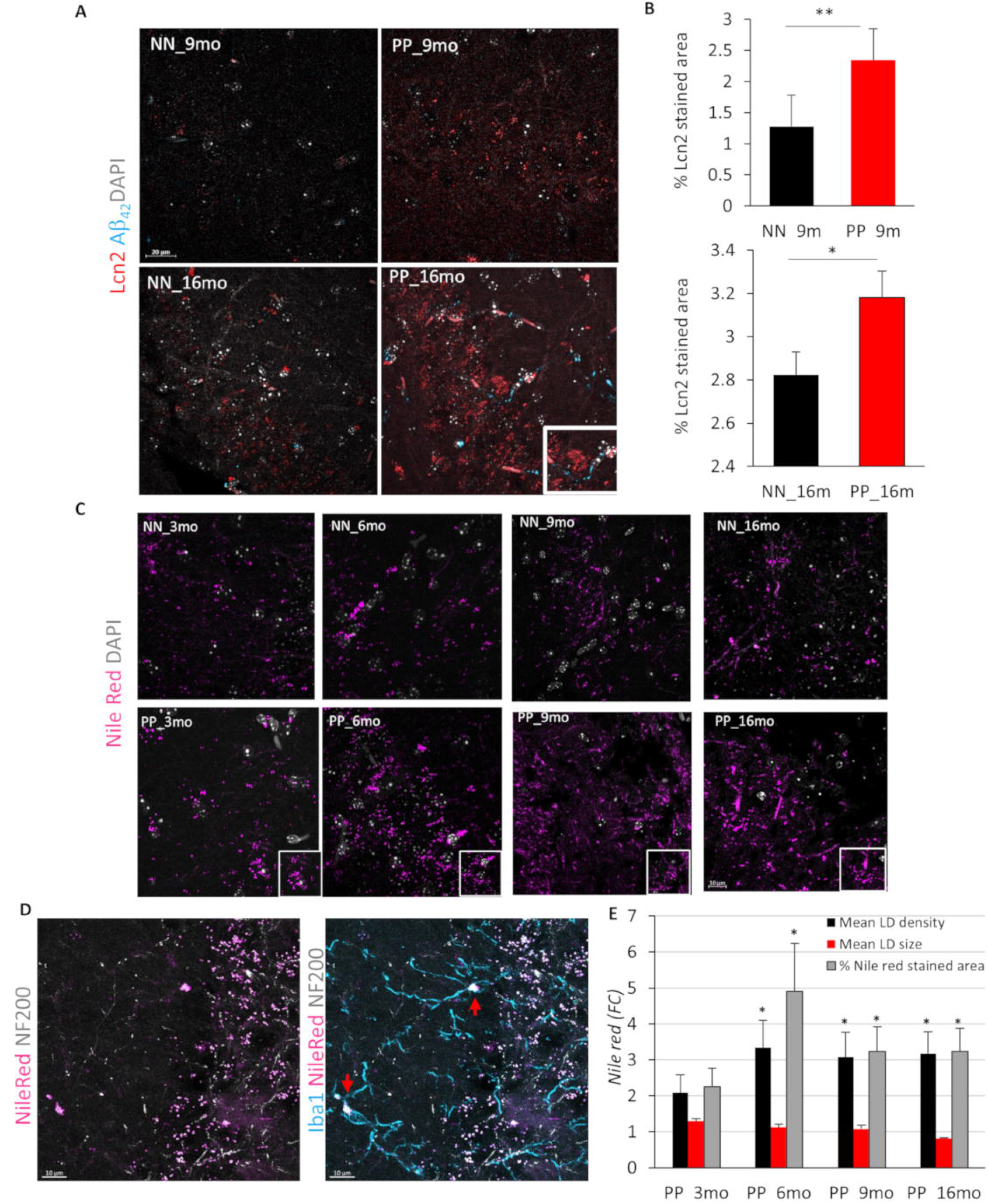
Effect of PolyI:C on the hippocampal Lcn2 & lipid droplet density. **(A)** Representative Lcn2, Aβ_1-42_ stained confocal images of saline and PolyI:C treated mice in 9 months and 16 months aged PP mice. Larger aggregates of Lcn2 (insert) visible at the 16 months PolyI:C exposed mice. **(B)** Quantification of % stained area indicates increased Lcn2 expression in the hippocampal CA3 field (n=4-6 mice for 9- and 16-months groups/treatment). **(C)** Nile red positive lipid droplet staining (violet) across aging in NN & PP mice. **(D)** Representative triple fluorescence labeling with Nile Red, Iba1 (teal) and NF200 (grey) shows an overlap between the different marker localized lipid droplets in Iba1 positive cells. Arrows indicating NileRed^+^/Iba1^+^ cells. **(E)** Bar chart representing fold change values in the lipid droplet density, mean size, and % stained area per ROI measured. Data are represented as mean±SEM relative to control (n=3-4 mice per group/treatment). *p<0.05, **p<0.005, Student’s t-test;. Scale bar A= 20μm, C,E= 10μm.

### Translation into Alzheimer’s disease

To assess whether the newly designed PolyI:C model is reproducing genetic changes in human AD, we performed targeted fingerprinting focusing in a cross-sectional cohort of post-mortem entorhinal cortices containing the hippocampus from age-matched subjects with mild-moderate AD, severe AD and age-matched healthy controls (CTL) (Table 3 and Suppl. Table 1). We first examined cell-type-specific genes, *GFAP*, *Iba1,* and *MAP2*. Consistently, with our model we observe a progressive increase in *GFAP* expression from Moderate to Severe AD (Fig. 7A and Suppl. Table 8), while *Iba1* and *MAP2* remain unchanged (Fig. 7A and Suppl. Table 8). Next, we examined some of the relevant DEG with reported association with AD and divided them into functional categories. *Glpr2* and *Ide* belonging to the glucose metabolism with differential expression in a 6 months old PP adult, did not show any significant difference between the clinical groups and controls (Fig. 7A and Suppl. Table 8). We next examined *Kcnj2* and *Egr2* which show downregulation in the PP model at 9 months (Table 2 and Fig. 5H) and observed high variability with no changes across stages (Suppl. Table 8 and Fig 7C). Among the cellular signaling genes, with specific repression in the PP model, we detect an opposite increasing trend in *c-Fos*, *c-Jun*, and *Notch1* to the PP model with a significant 3.1 upregulation of *c-Fos* in severe AD as compared to controls (Suppl. Table 8 and Fig 7D). In the lipid metabolism group, *Plin4* and *Lcn2* show no change opposite to the PP model (Suppl. Table 8 and Fig 7E), suggesting that those molecules are more implicated in vascular inflammation. To validate this assumption, we analyzed the vascular genes, *Klf4*, *Angptl4*, *Cyp1b1*, which showed transient alterations in the PP mode, and observed no change across stages (Suppl. Table 8 and Fig 7F). To understand the dependencies between the examined genes, we performed correlation analysis using the aggregate population. In general, all interactions are moderately significant (r>0.5) with positive associations between *c-fos* and *GFAP,* matching the increasing trend of the two transcripts, which suggest a cell-type-specific change, while *c-fos* is negatively associated with *MAP2* (Fig. 7G). *Kcnj2* is inversely associated with *Ide* reflecting their opposite trend, while *Ide* is positively correlated with *Cyp1b1* (Fig. 7G), indicating potential dependencies. Other interactions among the studied genes are seen but too subtle to be of functional relevance. Overall, despite the PolyI:C model replicates some aspects of the AD proteinopathy and microglia changes, the genetic expression between the mouse and humans differs substantially, raising the possibility of a mixed-vascular-AD model with diverse genetic fingerprints.

**Table 3.** Summary of Patients’ cohort I

| Stage | N. patients | F: M | Age (mean±SD) | Braak (mean ±SD) |
| --- | --- | --- | --- | --- |
| CTL | 9 | 5: 4 | 81± 8.8 | 1.0± 0.5 |
| MOD | 5 | 3: 2 | 85± 2.2 | 4.0± 1.6 |
| AD | 10 | 4: 6 | 82± 9.9 | 5.6± 0.5 |

CTL=healthy controls; MOD=moderate AD; AD=severe AD

**Figure 7.**
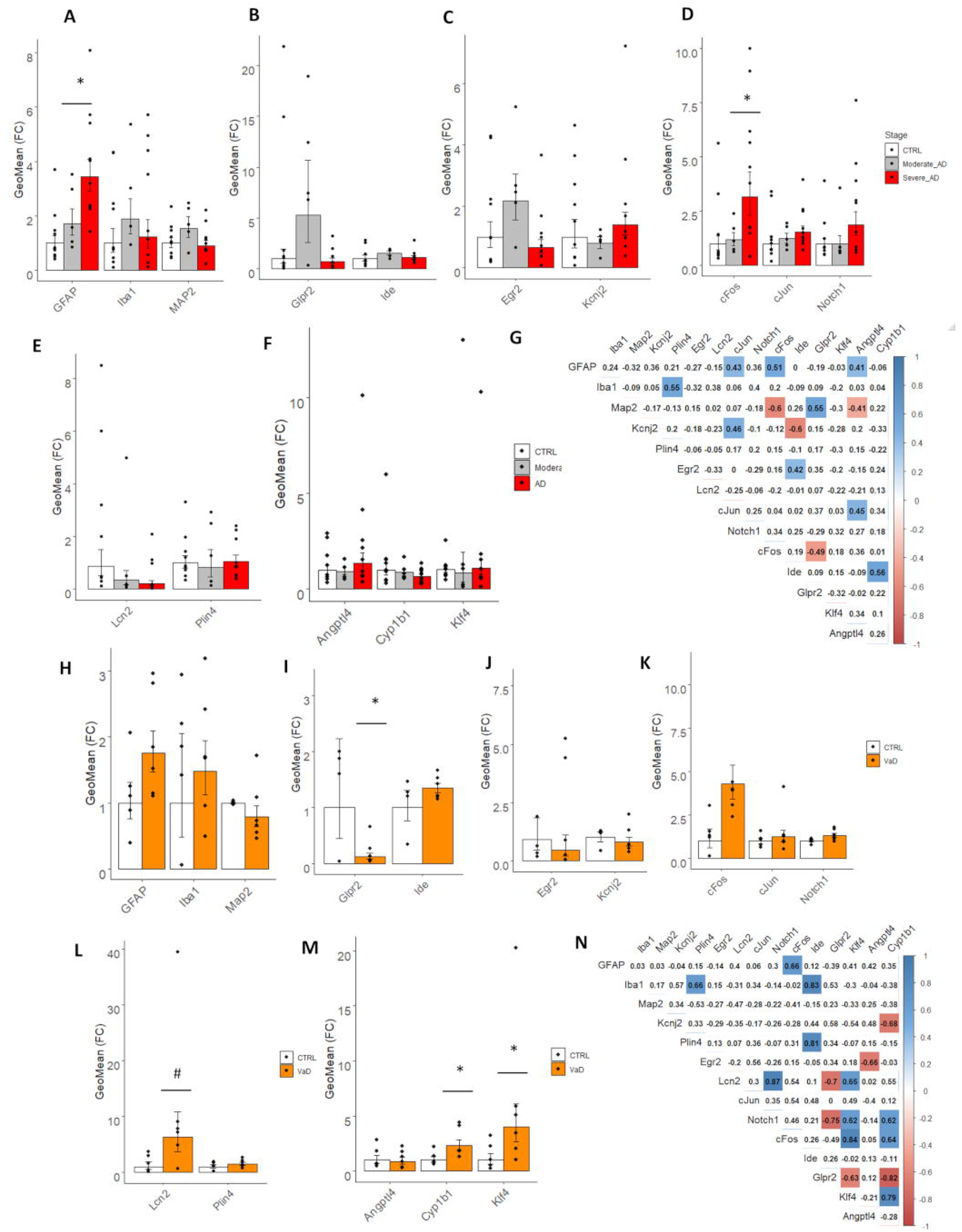
Target transcripts validation in the human brain specimen of patients with AD and VaD. Bar plots with Jitter showing the fold changes (FC) of the DEGs **(A-F)** in moderate AD and severe AD entorhinal cortices as compared to healthy controls (CTRL) and (**H-N)** in hippocampal sections from VaD as compared to healthy controls (CTRL). **(A and H)** FC of cell-type specific markers for astroglia (*GFAP*), microglia (*Iba1*) and neurons (*MAP2*). **(B and I)** FC of glucose metabolism markers, *Glpr2* and *Ide*. **(C and J)** FC of plasticity markers, *Egr2* and *Kcnj2*. (D and K) FC of cell signaling markers, *c-fos*, *c-Jun* and *Notch1*. **(E and L)** FC of inflammatory and metabolic markers, *Lcn2* and *Plin4*. **(F and M)**. FC of vascular markers, *Angptl4*, *Cyp1b1* and *Klf4*. **(G and N)** Correlation matrix of the selected biomarkers in the two study cohorts. Data are represented as Geometric mean of fold change± Geometric SEM relative to healthy controls. AD= Alzheimer’s disease and VaD=vascular dementia.

### Translation into vascular dementia

To verify the hypothesis of a mixed vascular-AD phenotype, we performed gene targets’ validation on a second cohort of hippocampi from vascular dementia patients and age-matched healthy controls (Table 4 and Suppl. Table 2). Cell type specific genes, *GFAP* and *Iba1*, indicate an increasing but not significant trend of microglia and astroglia, while MAP2 levels remain unchanged. Accordingly to our model, *Glpr2* decreases by 90% in vascular dementia as compared to controls (Fig. 7J and Supplementary Table 9) suggesting ongoing vascular hypoxia ((Xie et al. 2018)). On the other hand, the inflammatory and vascular markers, *Lcn2*, *Cyp1b1,* increase in the vascular dementia as in the PP model (Fig. 7L and 7K and Supplementary Table 9). Opposite to the genetic expression in the mouse, *Klf4*, *Notch1* and *c-fos* levels rise in vascular dementia (Fig. 7M and 7Y and Supplementary Table 9). Correlation analysis of the differentially expressed genes in the aggregate cohort indicates a positive association (r>0.6) among *Lcn2, Notch1 and Klf4*, implicating those factors in the vascular pathology. Whereas, along the mechanistic trajectory of vascular remodeling a negative association is observed between *Glpr2* and the two angiogenesis genes, *Klf4* and *Cyp1b1*. These results indicate that a handful of the gene targets that are significantly affected upon systemic inflammation in the brain of PP mice are reproduced in vascular neurodegenerative dementia.

**Table 4.**
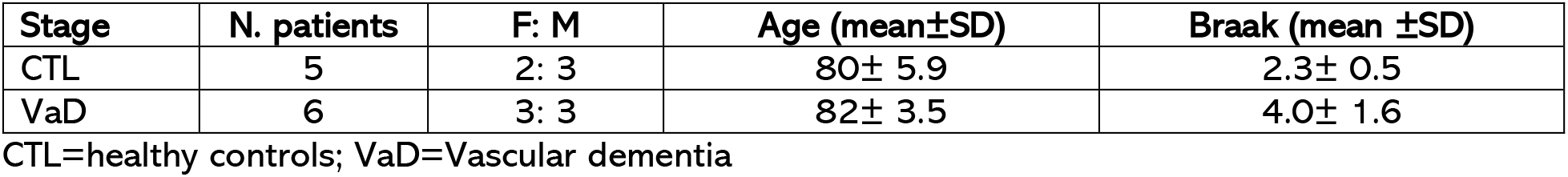
Summary of Patients’ cohort II

| Stage | N. patients | F: M | Age (mean±SD) | Braak (mean ±SD) |
| --- | --- | --- | --- | --- |
| CTL | 5 | 2: 3 | 80± 5.9 | 2.3± 0.5 |
| VaD | 6 | 3: 3 | 82± 3.5 | 4.0± 1.6 |

CTL=healthy controls; VaD=Vascular dementia

## Discussion

### Spread of peripheral inflammation to the brain

AD is a multifactorial complex disorder requiring the understanding of causal and risk factors for proper treatment. Currently, much research has been focused on the amyloid hypothesis. However, the large majority of clinical trials targeting amyloid-β have proved ineffective. Furthermore, evidence indicates that elderly people with Aβ accumulation in the brain may still preserve intact cognition questioning the role of amyloid-β in AD. On the other hand, modification of tau protein and its accumulation correlate better with the pathogenesis and topology of AD pathology (Joie et al. 2020). Nevertheless, if tau matches the neurodegenerative process, it remains a consequential event, while the etiology of sporadic AD lies mainly from the interplay of environmental and endogenous factors. There are increasing reports about a link between infectious diseases causing low-grade inflammatory responses over a lifetime and the development of AD with aging. Herpesviruses (Jamieson et al. 1991; Readhead et al. 2018), as well as bacteria such as Chlamydia Pneumoniae, Borrelia Burgdorferi and Porphyromonas Gingivalis (Hammond et al. 2010; Miklossy et al. 2004; Dominy et al. 2019), are enriched in the brains of AD patients likely playing a role in the disease pathogenesis. Experiments in transgenic mice for APP and Tau animal using viral, PolyI:C and bacterial, Lipopolysaccharides (LPS), aggravate and accelerate the AD-like pathology, supporting a role for infectious agents in triggering neuroinflammation and the proteinopathy (Krstic et al. 2012; Kitazawa et al. 2005). This prompted the development of a sporadic AD model using PolyI:C in naive non-transgenic mice to address whether a systemic sterile infection is a sufficient driver of the AD-pathology. This was first shown by the work of Kristic and colleagues demonstrated that a single prenatal immune challenge with PolyI:C could trigger an AD-like pathology and sustained brain inflammation (Krstic et al. 2012). Nevertheless, later studies showed no signs of long-term changes in central inflammatory responses or microglial morphology changes after a single prenatal immune activation (Giovanoli et al. 2015). Taking into consideration that infections typically occur over a lifetime, including the prenatal stages, we investigated whether systemic immune challenge using PolyI:C during prenatal (Wischhof et al. 2015; Arrode-Brusés and Brusés 2012; Patrich et al. 2016) and postnatal stages can induce a sustained peripheral inflammation with long-term effects on brain function mimicking the progressive pathophysiology of sporadic AD. The current work is the first cross-sectional and multi-modal examination of such a model, which is a prerequisite to mapping dynamic molecular changes occurring in the brain as a result of chronic peripheral inflammation. With the current experimental paradigm, we found that systemic infection with PolyI:C is associated with upregulation of inflammatory humoral factors (MCP-1, IL-6, IL-10, and TNF-α) in the plasma starting from 3 months age, with a peak at 6 months, which later decline in the aged mice until complete dissipation at 16 months. Interestingly, elevated IL-6 and IL-10 levels in the blood or brain of AD patients have been associated with the severity of cognitive decline and increased ventricular volume (Licastro et al. 2003; Leung et al. 2013) supporting the association between peripheral inflammation and AD. While the innate immune response to PolyI:C is observed at 3 and 6 months of age in the circulation, the adaptive immunity does not show any deviation (data not shown) and there is no observable infiltration of neutrophils, monocytes nor polymorphonuclear cells into the brain at these stages, suggesting preservation of the blood-brain barrier in younger animals until mid-age. Nevertheless, analysis of pro-inflammatory markers in the brain shows increased levels of IFN-γ at 6 and 9 months, while a rise in IL-6 at 6 months, precedes IL-1*β* increase at 9 months. This indicates that despite no cellular extravasation, cytokines at the earliest stages can spread from the periphery to the brain, causing and likely propagating neuroinflammation. On the other hand, in the oldest cohort, brain inflammation is exhausted, with an interesting inversion of IL-6 in PP mice suggesting immunodepression in the very aged also reflected by low levels of inflammatory cytokines in the circulation. Furthermore, it is known that IL-6 regulates the balance between pro- and anti-inflammatory responses depending on the context contributing to metabolic brain function (Borovcanin et al. 2017). For instance, IL-6^−/−^ mice develop obesity and insulin resistance indicating that deficiency of IL-6 can lead to metabolic changes (Wallenius et al. 2002). In parallel to the observed rise of neuroinflammation with aging, we also see an increase in microglia and astroglia gene expression at 6 and 9 months, reflecting cell-type-specific changes in these populations (Fig 1).

### Progressive proteinopathy

Along with the peripheral-central inflammatory responses, we observe a rise in hippocampal tau phosphorylation and p-tau/total tau ratio in accord with studies showing that IL-6 treatment induces tau hyperphosphorylation (p-Tau205/tau) in rat embryonic hippocampal neurons (Quintanilla 2004). In the aged PolyI:C mice, the hippocampal CA neurons accumulate phosphorylated tau protein in their axons with dominating astrogliosis. On the other hand, we observe that soluble A*β*_1-42_ increases at 6 months, coinciding with the rise in neuroinflammation and the onset of the tauopathy. Recent reports have indicated that inflammatory cytokines can increase *β*-secretase activity in neurons, producing elevated A*β*_1-42_ (Alasmari et al. 2018; Hur et al. 2020), which is perfectly in line with our model. This supports the use of anti-inflammatory agents at the early stages as a preventive strategy to the proteinopathy as recently reinforced (Hampel et al. 2020). In the aged PP mice, at 16 months, insoluble tangles or amyloid-*β* aggregates are visible in the hippocampus as small neuropil aggregates internalized at times by astroglia cells and in vessels reflecting a CAA. Interestingly, amyloid-*β* deposits in vessels are commonly seen in severe AD patients (53% of cases) but also in aged cognitively healthy individuals (50%) (Kövari et al. 2013). In our model, CAA-ike structures were however seen only in PP animals suggesting that the proteinopathy associated with vascular lesions is caused by systemic inflammation only. In the entorhinal cortex, gateway between the neocortex and the hippocampus controlling memory and spatial navigation (Fyhn et al. 2004), old PP mice display fibrillar aggregates. All these progressive changes are associated with working memory deficits displayed as a reduction in alternation at the Y-maze in agreement with the early prenatal and late postnatal PolyI:C mouse model (Krstic et al. 2012). Thus, spatial memory deficits upon PolyI:C challenge may be contributed by the progressive tauopathy in the limbic system accompanied by growing fibrillary aggregates disrupting the neural networks’ integrity responsible for spatial memory encoding.

### Microglia phenotypic change

Microglia are the major innate immune cells of the central nervous system mediating host defense responses against infectious agents, injury, abnormal accumulation of amyloid-*β*, and prion proteins (Yin et al. 2017). These cells express TLR3 viral receptors recognizing double-stranded RNA viruses (Town et al. 2006) and therefore play an important role in neuroinflammation in response to such stimuli initiating neuronal death. We have reported here a full-length characterization of microglial morphological changes across aging within the hippocampus of PolyI:C mice. Our results indicate dynamic changes in microglia soma features as well as its cell processes. Both pathological and non-pathological insults activate microglia changing from its normal ramified, rounded, and small soma to more hyper-ramified, reactive phagocytic morphology with aging (Walkera, Nilsson, and Jones 2013), which are observed in our model and are accompanied by the dynamic raise in inflammatory cytokines (IL-6, IL-10 and IFN-γ). In alignment with our study, others have reported an enlargement in the microglia soma/volume and an increase in their branch points after bacterial LPS exposure (Siemsen et al. 2020). As reviewed in other studies (Knuesel et al. 2014) the effect of second immune challenge postnatally exacerbates the AD-like pathology and the dynamic changes in the microglial morphology indicate a potential priming effect due to both maternal and early postnatal immune activation. Many models have been proposed to understand the morphological dynamic changes, for example after injury (Walker et al. 2014). In our study morphological changes in the aging PolyI:C mice resemble a microglial phenotype upon injury (Streit, Walter, and Pennell 1999) and after acute inflammatory response with neuraminidase treatment (Fernández-Arjona et al. 2017). While recently, it has been pointed out that the genetic repertoire in rodents and human microglia differs substantially (Masuda et al. 2019), the phenotypic transitions of microglia cells in this and other models recapitulates the dynamic undergoing changes in the progression of AD. Our and other findings support that neuroinflammation, passed on by the circulation and in response to the proteinopathy, is perpetuated influencing microglia cell fate to acquire a synapto- and neuro-toxic phenotype (Combs et al. 1999).

### Genetic remodeling

To further support the use of the PolyI:C mouse model as a viable preclinical experimental animal for AD research, we discover alterations in the hippocampal transcriptome relevant to inflammation and neurodegenerative dementia. GEA using the SynGO database indicates the biggest changes in presynaptic gene markers at the early stages (3 months), a steady-state at 6 months, and from 9 months on a progressive enrichment in post-synaptic genes. This is in line with the observed early presynaptic release of glutamate in response to oligomeric Amyloid-*β* (Palop and Mucke 2010), followed in time by post-synaptic scaling events aimed at preserving neuronal integrity at the expense of synaptic transmission (Findley et al. 2019). This mechanistic progression is theoretically confirmed by an aggregate GEA using KEGG, Wikipathway and Reactome pathways, which shows an early abundance in calcium signaling cascades, inflammation pathways, including MAPK signaling, PI3K-AKT signaling associated to cell survival and apoptosis. While at 6 months we confirmed an important decrease in *Glpr2* associated to synaptic depotentiation in response to hyperactivity (Sasaki-Hamada, Ikeda, and Oka 2019), (Xie et al. 2018; Bhusal et al. 2019), the changes in gene expression are much pronounced in the mid-aged 9 months and older 16 months PP mice with a 10% gene overlap. Longitudinal characterization revealed the upregulation of three genes (*Plin4, H2-Aa, Lcn2*) related to proinflammatory and glial function and downregulation of three genes (*Jun, Egr2, Kcnj2*) linked to neuronal or ion transport, reinforcing the mechanisms occurring in the progression of the neuropathology. We confirm a stark increase of *Lcn2* and *Plin4* at the late stages. *Lcn2* is a key gene involved in iron regulation and inflammation (Dekens et al. 2018). It is upregulated during systemic inflammation (Kang et al. 2017) and has been found to be elevated in the brains of AD patients (Naudé et al. 2012). *Plin4* is often associated with triacylglycerol metabolism and is involved in the biogenesis of lipid droplets in pathological degeneration (Han et al. 2018). The PolyI:C model shows a strong and specific rise in lipid droplets density accompanied by an increase in Lcn2 protein levels in neurons and glia, which confirms the metabolic and inflammatory disbalance in response to systemic immune challenge. The increase of Lcn2 in neurons and glial cells generates neuroinflammatory responses associated with insulin resistance and synaptic modulation (Song and Kim 2018)accumulation in the endothelial barrier (Ferreira et al. 2015) affects BBB permeability, as the genetic profiling in aging PP mice suggest. Thus the multifacet function of Lcn2 will result in increased neuroinflammation and reactive astrogliosis (Bi et al. 2013) which is confirmed in PP mice. The sterile-infection driven Lcn-2 supports its role as a putative druggable target to halt the immune-driven neuropathological progression.

### Reproducibility in postmortem tissue from AD and vascular dementia patients

One of the challenges of understanding the physiopathology of AD to develop targetable therapeutics is partially attributed to the poor reproducibility between animal models and humans and multifactorial etiology of the disease(Götz, Bodea, and Goedert 2018). Inflammatory, vascular processes and misfolded proteins should be considered as a whole and investigated closely to unravel dependencies. Furthermore, overlapping pathologies between AD and VaD, representing the most common forms of dementia, are present in more that 20% of the cases supporting common mechanisms and therapeutic investigations. Consistently, our study indicates that the PP model shows an AD proteinopathy coupled with vascular deficit. This conclusion derives from our DEG target repertoire analysis in two cohorts representing i) progressive AD stages and ii) vascular dementia. We confirm an astrogliosis in AD, captured by increased GFAP levels, comparable to the PP model while this effect is present but less pronounced in VaD. Astrogliosis has been shown to increase linearly with cognitive decline and is strongly associated with plaques and tangle formation (Serrano-Pozo et al. 2011). On the other hand in VaD, astrogliosis and astroglial endfeet swelling has been implicated as one of the triggering factors for vascular damage (Price et al. 2018; Wang et al. 2018). In both instances, reactive astrogliosis is a commonality of the two diseases associated with the production of pro-inflammatory cytokines such as IL-6, which is strongly upregulated in the PP model at 6 and 9 months. Along with the increase in *GFAP* expression, we see a positive association with *c-fos* and *Notch1* levels in the severe AD stage, supporting the proliferation of astroglia (Hisanaga et al. 1990) and the role of Notch1 in driving astroglia proliferation in response to inflammation through the proto-oncogene *c-fos* (Acaz-Fonseca et al. 2019). This data is in opposition to the PP mouse model, where despite an increase in astroglia, a decline in *c-fos* and *Notch1* is observed at 9 months. The discrepancy can be explained by the different signaling profiles of glia and neurons in rodents, suggesting that cellular cascades may be cell-specific depending on the species.

Investigating the genes involved in mediating vascular function (Deniz, Bozkurt, and Kurtel 2007), we observe no change in *Glpr2* expression in the progression of AD, while in VaD, *Glpr2* is downregulated similarly to the PP model supporting that cognitive deficit is contributed by lower blood perfusion ((Xie et al. 2018)). Along the same lines, the progressive increase in the inflammatory and metabolic markers, *Lcn2*, displayed by aging PP animals, is not reproduced in AD but in the VaD specimen. This finding is in contrast to the previous reports of a rise in Lcn2 protein levels in the hippocampus of severe AD subjects (Naudé et al. 2012) and patients with MCI (Choi, Lee, and Suk 2011), but is completely aligned with the recent reported upregulation of Lcn2 in CSF from VaD (n.d.; Llorens et al. 2020b). In support of the microvessel damage mediated by Lcn2 (J.-H. Kim et al. 2017), 9 months old PP mice report a rise in *Cyp1b1*, *Angptl4* and reduction in Klf4, which all regulate the BB permeability (Sangwung et al. 2017; Palenski et al. 2013; Huang et al. 2011). While in AD those markers remain unchanged, in VaD *Cyp1b1* is increased together with *Klf4*, while *Angptl4* remains unchanged. The different directionality of those vascular markers between mouse and human can be explained either by the time point of sampling or the species diversity. Nevertheless, a vascular pathology in the PP model is supported by two-fold reduction of Claudin 5, *Cldn5*, a key regulator of BBB permeability. While mixed pathologies in neurodegenerative diseases are not uncommon accounting for more than 20% of the cases (Custodio et al. 2017), the PP model shows a mixed pathology with pronounced neuroinflammation, AD-like proteinopathy, and vascular factors. Interestingly, a recent finding has shown that hyperphosphorylated Tau affects neurovascular coupling (Park et al. 2020) bridging AD-mechanism to the vascular deficit. We acknowledge that the number of human samples analyzed is small, however the translational validation of the PolyI:C data clearly demonstrates that this mouse model can reproduce some of the characteristic features of reactive central inflammation (Hampel et al. 2020) and vascular pathology encountered in neurodegenerative dementia.

## Conclusions

The present research, using the PolyI:C mouse model of sterile infection, demonstrates that chronic systemic inflammation during adulthood causes a progressive neuropathology with neuroinflammation, insoluble protein aggregates, vascular permeability, microglia remodeling and behavioral deficits, mimicking a mixed vascular-AD pathology. While the PolyI:C model has been previously employed in the prenatal stage or in old age, the present longitudinal study adds to the understanding of the effects of sustained inflammation on brain homeostasis over a lifetime. Our post-mortem analysis on AD and VaD brain specimens, shows partially overlapping genetic profiles between VaD and the PolyI:C mouse supporting that systemic inflammation can cause vascular deficit besides neuroinflammation and proteinopathy. Indeed, chronic inflammation is known to pose a risk for cardiovascular health which with aging may contribute to overlapping pathologies (Metti and Cauley 2012; Newcombe et al. 2018). The study is of use not only for the understanding of the interplay between peripheral and central processes but also presents a surrogate animal model displaying a sporadic vascular AD-mixed phenotype, which can be used for testing therapeutics against pathological brain aging.

## List of Abbreviations

ABCA7: ATP-binding cassette sub-family A member 7
AD: Alzheimer’s disease
Angptl4: Angiopoietin like 4
ApoE: Apolipoprotein E
ApoEε4: Apolipoprotein E variant ε4
Aβ: Amyloid beta
BBB: Blood Brain Barrier
BIN1: Box-dependent-interacting protein 1
CA: Cornu Ammonis
CAA: Cerebral Amyloid Angiopathy
CD2AP: CD2 Associated Protein
CD33: Myeloid Cell Surface Antigen CD3
CELF1: CUGBP Elav-like family member 1
c-fos: Proto-oncogene c-Fos
Cyp1b1: Cytochrome P450 1B1
Cldn5: Claudin 5
CLU: Clusterin
CNS: Central Nervous System
CR1: Complement receptor type 1
CSF: Cerebrospinal fluid
CTL: Control
Cx: Cortex
DABCO: 1,4-diazabicyclo [2.2.2] octane
DAPI: 4′,6-diamidino-2-phenylindole
DEGs: Differentially Expressed Genes
dsRNA: Double-strand RNA
EC: Entorhinal cortex
EDTA: Ethylenediaminetetraacetic acid
EGR2: Early growth response protein 2
EPHA1: Ephrin type-A receptor 1
FERMT2: Fermitin family homolog 2
GD: Gestational Day
GEA: Gene enrichment analysis
GFAP: Glial fibrillary acidic protein
GlpR2: Glucagon like peptide 2 receptor
GO: Gene ontology
GRIN1: Glutamate Ionotropic Receptor NMDA Type Subunit 1
GWAS: Genome Wide Association Studies
H2-Aa: H-2 class II histocompatibility antigen, A-B alpha chain
Iba-1: Ionized calcium binding adaptor molecule1
Ide: Insulin degrading enzyme
IL: Interleukin
INPP5D: Phosphatidylinositol 3,4,5-trisphosphate 5-phosphatase 1
KCNJ2: Potassium Inwardly Rectifying Channel Subfamily J Member 2
KEGG: Kyoto Encyclopedia of Genes and Genomes
Klf4: zinc finger-containing Krüppel-like factor
KS: Kolmogorov-Smirnov
c-Jun: Transcription factor AP-1
LCN2: Lipocalin 2
LPS: Lipopolysaccharide
MAP2: Microtubule-associated protein 2
MAPK: Mitogen-activated protein kinase 1
MCI: Mild Cognitively Impairment
MCP-1: Monocyte Chemoattractant Protein-1
MEF2C: Myocyte-specific enhancer factor 2C
mM: milliMolar
M-MLV: Moloney Murine Leukemia Virus Reverse Transcriptase mo month
mRNA: Messenger RNA
MS4A4A: Membrane-spanning 4-domains subfamily A member 4A
ND: Neurodegenerative disease
NME8: Thioredoxin domain-containing protein 3
NN: Saline controls
Notch1: Neurogenic locus notch homolog protein 1
PFA: Paraformaldehyde
PICALM: Phosphatidylinositol-binding clathrin assembly protein
PLCG2: 1-phosphatidylinositol 4,5-bisphosphate phosphodiesterase gamma-2
pM: picomolar
PLIN4: Perilipin 4
PMNs: Polymorphonuclear neutrophils
PolyI:C: Polyinosinic:Polycytidylic acid
PSEN: Presenilin
PP: PolyI:C injected animals
pTau: Phosphorylated tau
PVA: Polyvinyl alcohol
rpm: revolutions per minute
rRNA: ribosomal RNA
r: Spearman’s Rank Correlation
SEM: Standard Error Mean
SLC24A4: Sodium/potassium/calcium exchanger 4
SORL1: Sortilin-related receptor
SRCAP: Snf2-related CREBBP activator protein
TBS: Trizma-based salt solution
TLRs: Toll Like receptors
TNFα: Tumor necrosis factor alpha
TREM2: Triggering receptor expressed on myeloid cells 2
ZCWPW1: Zinc finger CW-type PWWP domain protein 1
μg: microgram
μm: micrometer

## Declarations

### Ethical Approval and Consent to participate

Animal experimentation was approved by the animal experiment committee, University of Fribourg (Protocol no. 2016_32_FR registered 01/01/2017).

The use of human tissue has been approved by the Ethical Commission of the Brain Bank for Dementia UK (OBB443 registered 1/05/2017 and OB344 registered 1/02/2014), Stanford (Stanford IRB), and the Ethical Commission from the Canton of Fribourg and Vaud (N. 325/14). All experiments conducted on human tissue comply with the WMA Declaration of Helsinki.

### Consent for publication

All authors agree on publishing the original data

### Availability of supporting data

Supporting data is available in the form of supplementary material and tables. All raw data is available on request.

### Authors’ contributions

PB conducted the main bulk of the experiments and contributed to the writing. ID performed the bioinformatic analysis. EZ and ET performed the cellular immunology experiments from blood and brain. AF and EB performed the blind quantitative analysis of microglia morphology using custom-made scripts. MAD assisted in the inflammatory model characterization. LA designed the study and wrote the manuscript. All authors read and approved the final manuscript

## Acknowledgments

We would like to thank Mrs. E. Martin and V. Tache for the technical support. We are also grateful to Dr. M. Reggente in providing his expertise in R. We thank Prof. T. Montine for sharing the brain tissue from the Stanford Biobank for this study. We are thankful to the donors and their families for allowing us access to the human tissue (Oxford, UK and Stanford, USA).

