## Supplementary material for "Systemic inflammation causes microglial dysfunction with a mixed AD-like pathology": Suppl. Tables

**Supplementary Tables**

**Supplementary Table 1.** Procured Human tissues of cohort I

| **UNIP** | **Age** | **Gender** | **Diagnosis*** | **Braak** | **Plaques** | **Condition** | **Biobank** |
| --- | --- | --- | --- | --- | --- | --- | --- |
| NP13/074 | 74 | Female | Not AD | I | No plaques | CTRL | Oxford |
| NP12/132 | 67 | Female | Not AD | I | No plaques | CTRL | Oxford |
| LA22 | 91 | Male | Not AD | I (B1) | No neuritic plaques (C0) | CTRL | Stanford |
| LA33 | 89 | Female | Not AD | II (B1) | No neuritic plaques (C0) | CTRL | Stanford |
| NP 169/11 | 84 | Female | Not AD | I | No plaques | CTRL | Oxford |
| NP 071/13 | 83 | Female | Not AD | I | No plaques | CTRL | Oxford |
| NP 121/12 | 89 | Female | Not AD | I | No plaques | CTRL | Oxford |
| NP13/081_ | 70 | Male | Not AD | I | No plaques | CTRL | Oxford |
| RI03/211 | 85 | Male | Not AD | 0 | No plaques | CTRL | Oxford |
| NP13/163 | 89 | Female | High | IV | Sparse | Moderate | Oxford |
| NP15/099 | 83 | Male | High | IV | Moderate (C2) | Moderate | Oxford |
| RI04/43 | 85 | Female | Low | III | No plaques | Moderate | Oxford |
| NP13/160 | 88 | Female | High | IV | Not available | Moderate | Oxford |
| RI05/14 | 87 | Male | High | IV | Sparse plaques | Moderate | Oxford |
| LA66 | 95 | Male | High | VI (B3) | Frequent (C3) | AD | Stanford |
| LA88 | 84 | Male | High | VI (B3) | Frequent (C3) | AD | Stanford |
| AP-J10 | 76 | Male | Not assessed | V (B3) | Frequent (C3) | AD | Stanford |
| AP-H8 | 94 | Male | High | VI (B3) | Frequent (C3) | AD | Stanford |
| NP 133/12 | 78 | Male | Definite AD | V | Neuritic plaques | AD | Oxford |
| NP 035/13 | 85 | Male | Definite AD | V | Neuritic plaques | AD | Oxford |
| LA77 | 87 | Female | High | VI (B3) | Frequent (C3) | AD | Oxford |
| NP128/14 | 61 | Female | Definite AD | VI | Neuritic plaques | AD | Oxford |
| NP14/033 | 77 | Female | Definite AD | V | Neuritic plaques | AD | Oxford |
| AP-G7 | 85 | Female | Not assessed | VI (B3) | Moderate (C2) | AD | Stanford |

*= probability of AD case; CTL=healthy controls; MOD=moderate AD; AD=severe AD

**Supplementary Table 2.** Procured Human tissues of cohort II

| **UNIP** | **Age** | **Gender** | **Diagnosis** | **Braak** | **Condition** | **Biobank** |
| --- | --- | --- | --- | --- | --- | --- |
| 2017-016 | 88 | Male | No dementia | I | CTRL | Netherlands Brain Biobank |
| 2017-078 | 88 | Female | No dementia | III | CTRL | Netherlands Brain Biobank |
| 2017-109 | 91 | Male | No dementia | II | CTRL | Netherlands Brain Biobank |
| 2017-131 | 71 | Female | No dementia | II | CTRL | Netherlands Brain Biobank |
| 2018-028 | 77 | Female | No dementia | II | CTRL | Netherlands Brain Biobank |
| 1998-073 | 86 | Female | Vasc. dementia | III | VaD | Netherlands Brain Biobank |
| 2007-043 | 78 | Male | Vasc. dementia | III | VaD | Netherlands Brain Biobank |
| 2008-060 | 84 | Male | Vasc. dementia | III | VaD | Netherlands Brain Biobank |
| 2010-086 | 82 | Female | Vasc. dementia | III | VaD | Netherlands Brain Biobank |
| 2010-092 | 85 | Male | Vasc. dementia | III | VaD | Netherlands Brain Biobank |
| 2010-098 | 78 | Female | Vasc. dementia | III | VaD | Netherlands Brain Biobank |

VaD= vascular dementia

**Supplementary Table 3.** Primers for qPCR

| **Gene** | **Forward** | **Reverse** | **Species** |
| --- | --- | --- | --- |
| *IL-6* | 5'ATG GAT GCT ACC AAA CTG GAT3' | 5'TGA AGG ACT CTG GCT TTG TCT3' | Mouse |
| *GLPR2* | 5'GGT CCT CCT GCA CTA CTT T3' | 5'CCA GGG AAT AAC AAA CAG C3' | Mouse |
| *IDE* | 5'CCA GGT AGT GGG GTG GTT TT3' | 5'TAA TGG GCT ATG TGC GCG TG3' | Mouse |
| *Kcnj2* | 5'CAC AGC TTC TCA AAT CTA GGA TCA3' | 5'CTA TTT CGT GAA CGA TAG TGA TGG3' | Mouse |
| *c-Jun* | 5'ACG ACC TTC TAC GAC GAT GC3' | 5'CCA GGT TCA AGG TCA TGC TC3' | Mouse |
| *c-fos* | 5'CGG GTT TCA ACG CCG ACT A3' | 5'TGG CAC TAG AGA CGG ACA GAT3' | Mouse |
| *Notch1* | 5'TCA GAG GCC AGC AAG AAG AA3' | 5'GCT CCT CAA ACC GGA ACT TC3' | Mouse |
| *Egr2* | 5'CCG CCA AGG CCG TAG ACA AAA3' | 5'GGG TCA ATG GAG AAC TTG CCC3' | Mouse |
| *Lcn2* | 5'TTT CAC CCG CTT TGC CAA GT3' | 5'GTC TCT GCG CAT CCC AGT CA3' | Mouse |
| *PLIN4* | 5'GAC CAG CAG TGA AGA TGC CT3' | 5'TCC TTC GTA TTG GTG AGG AC3' | Mouse |
| *Klf4* | 5'CGA CTA ACC GTT GGC GTG A3' | 5'TGG GTT AGC GAG TTG GAA AGG3' | Mouse |
| *Cyp1b1* | 5'GGA CAA GGA CGG CTT CAT TA3' | 5'GCG AGG ATG GAG ATG AAG AG3' | Mouse |
| *Angptl4* | 5'GGG ACC TTA ACT GTG CCA AG3' | 5'GAA TGG CTA CAG GTA CCA AAC C3' | Mouse |
| *β-actin* | 5'GTG ACG TTG ACA TCC GTA AAG A3' | 5'GCC GGA CTC ATC GTA CTC C3' | Mouse |
| *GFAP* | 5'CCT CTC CCT GGC TCG AAT G3' | 5'GGA AGC GAA CCT TCT CGA TGT A3' | Human |
| *Iba1* | 5'CTC AGG ATG ATG CTG GGC AAG AGA3' | 5'AGC CCC TTC AAT CCC ATC ATC CCT3' | Human |
| *MAP2* | 5'CTG CTT TAC AGG GTA GCA CAA3' | 5'TTG AGT ATG GCA AAC GGT CTG3' | Human |
| *GLPR2* | 5'ACC TTG GTG GAG TGA AGA GAG3' | 5'GCC AAA TAT CCG TGG CGT TC3' | Human |
| *IDE* | 5'GCC GAA GGC TTG TCT CAA CT3' | 5'CAA ATA GGC CAT GTT ACA GTG CAA3' | Human |
| *Kcnj2* | 5'TGG ATG CTG GTT ATC TTC TGC3' | 5'AGC CTA TGG TTG TCT GGG TCT3' | Human |
| *c-Jun* | 5'GAG CTG GAG CGC CTG ATA AT3' | 5'CCC TCC TGC TCA TCT GTC AC3' | Human |
| *c-fos* | 5'AGG AGG GAG CTG ACT GAT ACA CT3' | 5'TTT CCT TCT CCT TCA GCA GGT T3' | Human |
| *Notch1* | 5'GAG GCG TGG CAG ACT ATG C3' | 5'CTT GTA CTC CGT CAG CGT GA3' | Human |
| *Egr2* | 5'CCG CCA AGG CCG TAG ACA AAA3' | 5'GGG TCA ATG GAG AAC TTG CCC3' | Human |
| *Lcn2* | 5'TCA CCC TCT ACG GGA GAA CC3' | 5'GGG ACA GGG AAG ACG ATG TG3' | Human |
| *PLIN4* | 5'CTG GTG GCC AAC GCA CAT AG3' | 5'GCC CCG GAC ACC ATC TTT TC3' | Human |
| *Klf4* | 5'ACC TTC TTC ACC CCT AGA GC3' | 5'AAGGTTTCTCACCTGTGTGG3' | Human |
| *Cyp1b1* | 5'TCC TCC TCT TCA CCA GGT ATC C3' | 5'GGC TGG TCA CCC ATA CAA GG3' | Human |
| *Angptl4* | 5'ATG GCT CAG TGG ACT TCA AC3' | 5'GCT ATG CAC CTT CTC CAG AC3' | Human |
| *β-actin* | 5'CGC GAG AAG ATG ACC CAG AT3' | 5'GAT AGC ACA GCC TGG ATA GCA AC3' | Human |
| *hRPL13A* | 5'AAA AGC GGA TGG TGG TTC CT3' | 5'GCT GTC ACT GCC TGG TAC TT3' | Human |

**Supplementary Table 4 .** Locomotor activity in PP and NN mice

| **Age** | **Group** | **No. of arm entries**  **(Y maze spontaneous alternation task)** |
| --- | --- | --- |
| 3 months | NN | 18.8±1.1 |
|  | PP | 21.6±2.5 |
| 6 months | NN | 16.6±1.2 |
|  | PP | 17.5±1.7 |
| 9 months | NN | 11±0.9 |
|  | PP | 10.9±0.8 |
| 16 months | NN | 10±0.7 |
|  | PP | 11±1.1 |

Data are represented as mean ± SEM. NN= prenatal& postnatal saline injected controls, PP = prenatal and postnatal PolyI:C treated mice. 3 months (NN=8, PP=9), 6 months (NN=8, PP=8), 9 & 16 months (NN, PP=8 each).

**Supplementary Table 5** Anxiety testing in aging PP and NN

| **Age** | **Group** | **Anxiety Testing** | | |
| --- | --- | --- | --- | --- |
|  |  | **Elevated O-maze**  **(% time in open arm)** | **Light/Dark test**  **(% time in the light)** | **Open Field test**  **(% time spent in center)** |
| 3 months | NN | 10.2±1.7 | 31.2±4.0 | 23.4±10.0 |
|  | PP | 7.1±4.1 | 40.8±4.0 | 20.4±5.9 |
| 6 months | NN | 7.1±3.1 | 23.8±27.2 | 32.5±20.9 |
|  | PP | 9.0±4.6 | 22.3±9.1 | 39.2±11.2 |
| 9 months | NN | - | - | 21.1±3.7 |
|  | PP | - | - | 25.7±5.1 |
| 16 months | NN | - | - | 4.4±1.25 |
|  | PP | - | - | 6.8±3.0 |

Data are represented as mean ± SEM. NN= prenatal& postnatal saline injected controls, PP = prenatal and postnatal PolyI:C treated mice. 3 months (NN=8, PP=9), 6 months (NN=8, PP=8), 9 & 16 months (NN, PP=8 each).

**Supplementary Table 6.** Quantification of Iba-1 positive cells per volume

| **Age** | **Group** | **Iba-1 + cells/ mm^3^** |
| --- | --- | --- |
| 3 months | NN | 7226±292 |
|  | PP | 7567±341 |
| 6 months | NN | 8792±420 |
|  | PP | 8806±379 |
| 9 months | NN | 8111±362 |
|  | PP | 7884±342 |
| 16 months | NN | 7433±460 |
|  | PP | 7016±411 |

**Supplementary Table 7.** GO Analysis of the top 10 significant Biological Processes for DEGs between PP and NN

| **Age** | **GO BP** | **% Ass. Genes** | **Upregulated**  **(in descending order)** | **Downregulated**  **(in ascending order)** |
| --- | --- | --- | --- | --- |
| **3 months** | positive regulation of synaptic transmission | 2.13 | Calb2- | [Cckbr, Cux2, Camk2d, Tshz3] |
|  | extracellular matrix organization | 1.69 | Col8a1 | [Lamc2, Rxfp1, Fn1] |
|  | regulation of cytosolic calcium ion concentration | 1.33 | [Trpv4, Calb2] | [Cckbr, Camk2d, Cacna1g, Mchr1] |
|  | secretion | 0.81 | [Prl, Trpv4, Trp73] | [Cplx3, Cckbr, Unc13c, Olfm2, Srebf1, Fn1, Cacna1g] |
| **6 months** | cell morphogenesis involved in differentiation | 0.71 | [Xlr3b] | [Col12a1, Lhx9, Trpc6] |
|  | regulation of nervous system development | 0.57 | [Xlr3b, H2-Q4, Aspa,] | [Lrtm2, Trpc6] |
| **9 months** | regulation of endothelial cell differentiation | 17.07 | - | [Apold1, Cldn5, Acvrl1, Id1, Kdr, Btg1, Notch1] |
|  | sprouting angiogenesis | 8.27 | [Adamts9, Rhoj] | [Klf4, Dll4, Klf2, Nr4a1, Nrarp, Flt1, Kdr, Sema4c, Notch1] |
|  | *regulation of neural precursor cell proliferation* | 7.03 | [H2-Aa] | [Btg2, Dll4, Gli2, Gli3, Nes, Notch1, Sox2, Sox21] |
|  | negative regulation of cell adhesion | 4.43 | [Trpv4, H2-Aa, Cyp1b1, Lrrc23, Dlg5] | [Klf4, Dusp1, Acer2, Nrarp, Cxcl12, Acvrl1, Gli3, Sema4c, Notch1] |
|  | morphogenesis of an epithelium | 4.03 | [Mgp, Fgfr2, Agt, Tgm2, Dlg5, Egfr] | [Cxcl12, Zic2, Sox21, Flt1, Sox2, Acvrl1, Gli3, Kdr, Spry2, Sema4c, Notch1] |
|  | *negative regulation of cell population proliferation* | 3.85 | [Fap, H2-Aa, Cyp1b1, Lrrc23, Inhba, Gjb6, Fgfr2, Agt, Cdkn1a, Dlg5] | [Btg2, Klf4, Dll4, Dusp1, Gkn3, Wnt9a, Ifit3, Jun, Plk5, Sox21, Flt1, Sox2, Krt9, Acvrl1, Gli3, Apln, Tob1, Spry2, Btg1, Notch1] |
|  | Response to peptide | 3.52 | Trpv4, Baiap2l1, Lepr, Ucp2, Sgk1, , Mc4r, Eprs, Agt, Fbn1, Klf15, Nfkbia] | [Btg2, Cd40, Cxcl12, Egr1, Egr2, Id1, Klf2, Klf4, Notch1, Nr4a1, Ucp2] |
|  | *positive regulation of cell death* | 2.93 | [Fap, Cyp1b1, Inhba, Ucp2, Sbno2, , Agt, Gadd45g, Cdkn1a, , Tgm2] | [Cd40, Btg2, Klf4, Klf2, Nr4a1, Egr2, Cxcl12, Egr1, Id1, Notch1] |
|  | regulation of response to external stimulus | 2.82 | [Pla2g5, Anxa2, Cmklr1, Pros1, Il1r1, Sbno2, Agt, Cdkn1a, Tgm2, Nfkbia, Egfr] | [Cxcl10, Klf4, Foxf1, Dusp1, Sema3g, Cxcl12, Zfp36, Krt9, Ctla2a, Kdr, Sema4c, Notch1] |
|  | *Response to cytokine* | 2.63 | [Pla2g5, H2-Aa, Lepr, H2-Eb1, Spp1, Osmr, Il1r1, Eprs, Sbno2, Nfkbia, Traip] | [Cd40, Cxcl10, Klf4, Foxf1, Fos, Dusp1, Gch1, Klf2, Ifit3, Junb, Jun, Cxcl12, Zfp36, Egr1, Spry2, C1qtnf4] |
| **16 months** | Response to metal ion | 2.04 | [Xdh, Mt2, Cp, Mt1, Txnip] | [Lct, Capn3, Abcc8, Jun] |
|  | *Axon development* | 1.80 | [Tsku, Cntn6, Apod, Cntn4] | [Egr2, Robo3, Lhx9, Jun, Auts2, Islr2] |
|  | regulation of hormone levels | 1.71 | [Esr1, Sfrp1, Adcyap1, Edn3, Sult1a1, Cplx3, Tac1] | [Iyd, Abcc8, Myt1, Neurod1] |
|  | negative regulation of cell population proliferation | 1.54 | [Esr1, Lrrc32, Sfrp1, Nupr1, Adcyap1, Xdh, Ifit3, Apod] | [Pla2g2f, Cblb, Abcc8, Jun] |
|  | *response to oxygen-containing compound* | 1.12 | [Esr1, Fbln5, Gjb2, Col1a2, Sfrp1, Acer2, Adcyap1, Tac1, Igfbp7, Apod, Mt1, Txnip] | [Lct, Tdo2, Egr2, Trpc6, Zbtb20, Abcc8, Myt1, Jun, Neurod1, Elk1] |

**Supplementary Table 8**. Investigation of target transcripts in entorhinal cortex from moderate and severe AD patients

| **Category** | **Gene** | **Stage** | **DCq (Mean±SEM)** | **FC** | **P value** | **Relation to Model** |
| --- | --- | --- | --- | --- | --- | --- |
| *Cell Type* | *GFAP* | *CTL* | 1.4± 0.4 | - |  | - |
|  |  | *MOD* | 0.6± 0.5 | 1.7 | 0.25 | - |
|  |  | *AD* | -0.4± 0.2 | 3.4 | 0.003* | consistent |
|  | *Iba1* | *CTL* | 1.3± 0.6 | - | - | - |
|  |  | *MOD* | 0.6± 0.5 | 1.9 | 0.19 | - |
|  |  | *AD* | -0.3± 0.7 | 1.2 | 0.75 | - |
|  | *Map2* | *CTL* | 8.6± 0.3 | - | - | - |
|  |  | *MOD* | 8.0± 0.4 | 1.5 | 0.19 | - |
|  |  | *AD* | 8.4± 0.3 | 0.9 | 0.75 | - |
| *Glucose metabolism* | *Glpr2* | *CTL* | 7.3± 1.0 | - | - | - |
|  |  | *MOD* | 4.8± 1.2 | 5.2 | 0.29 | opposite |
|  |  | *AD* | 7.7± 0.8 | 0.7 | 0.22 | consistent |
|  | *Ide* | *CTL* | 7.6± 0.4 | - | - | - |
|  |  | *MOD* | 7.0± 0.2 | 1.5 | 0.68 | - |
|  |  | *AD* | 7.5± 0.2 | 1.1 | 0.68 | - |
| *Synapse* | *Kcnj2* | *CTL* | 5.3± 0.7 | - | - | - |
|  |  | *MOD* | 5.5± 0.4 | 0.8 | 0.13 | consistent |
|  |  | *AD* | 4.7± 0.4 | 1.4 | 0.82 | - |
|  | *Egr2* | *CTL* | 11.5± 0.6 | - | - | - |
|  |  | *MOD* | 10.4± 0.5 | 2.1 | 0.34 | - |
|  |  | *AD* | 12.1± 0.5 | 0.7 | 0.28 | - |
| *Cellular signaling* | *c-Jun* | *CTL* | 4.9± 0.4 | - | - | - |
|  |  | *MOD* | 4.6± 0.3 | 1.2 | 0.90 | - |
|  |  | *AD* | 4.3± 0.2 | 1.5 | 0.46 | - |
|  | *c-Fos* | *CTL* | 4.2± 0.5 | - | - | - |
|  |  | *MOD* | 3.9± 0.4 | 1.2 | 0.71 | - |
|  |  | *AD* | 2.4± 0.5 | 3.1 | 0.03* | opposite |
|  | *Notch1* | *CTL* | 5.9± 0.3 | - | - | - |
|  |  | *MOD* | 5.9± 0.5 | 1.0 | 0.95 | - |
|  |  | *AD* | 5.2± 0.4 | 1.85 | 0.10 | opposite |
| *Lipid metabolism* | *PLIN4* | *CTL* | 8.2± 0.3 | - | - | - |
|  |  | *MOD* | 8.4± 0.9 | 0.8 | 0.78 | - |
|  |  | *AD* | 8.1± 0.3 | 1.0 | 0.99 | - |
|  | *Lcn2* | *CTL* | 8.5± 0.9 | - | - | - |
|  |  | *MOD* | 10.0± 1.2 | 0.3 | 0.41 | - |
|  |  | *AD* | 10.8± 0.7 | 0.2 | 0.10 | opposite- |
| *Infalmmatioon*  *Vascular remodeling* | *Klf4* | *CTL* | 11.4± 0.2 | - | - | - |
|  |  | *MOD* | 11.0± 0.8 | 0.8 | 0.84 | - |
|  |  | *AD* | 11.7± 1.2 | 1.2 | 0.65 | - |
|  | *Cyp1b1* | *CTL* | 5.7± 0.3 | - | - | - |
|  |  | *MOD* | 5.9± 0.3 | 0.9 | 0.73 | - |
|  |  | *AD* | 6.2± 0.3 | 0.7 | 0.26* | - |
|  | *Angptl4* | *CTL* | 4.5± 0.4 | - | - | - |
|  |  | *MOD* | 4.7± 0.4 | 0.9 | 0.71 | - |
|  |  | *AD* | 4.1± 0.5 | 1.3 | 0.49 | - |

**Supplementary Table 9**. Investigation of target transcripts in entorhinal cortex from control and Vascular dementia patients

| **Category** | **Gene** | **Stage** | **DCq (Mean±SEM)** | **FC** | **P value** | **Relation to Model** |
| --- | --- | --- | --- | --- | --- | --- |
| *Cell Type* | *GFAP* | *CTL* | 3.1± 0.9 | - |  | - |
|  |  | *VaD* | 2.3± 0.6 | 1.8 | 0.125* | consistent |
|  | *Iba1* | *CTL* | 11.5± 2.3 | - | - | - |
|  |  | *VaD* | 10.9± 1.0 | 1.5 | 0.63 | - |
|  | *MAP2* | *CTL* | 9.7± 0.3 | - | - | - |
|  |  | *VaD* | 10.0± 0.7 | 0.8 | 0.27 | - |
| *Glucose metabolism* | *Glpr2* | *CTL* | 8.7± 2.6 | - | - | - |
|  |  | *VaD* | 12.2± 1.1 | 0.09 | 0.04 | consistent |
|  | *Ide* | *CTL* | 10.4± 0.9 | - | - | - |
|  |  | *VaD* | 9.9± 0.2 | 1.3 | 0.33 | - |
| *Synapse* | *Kcnj2* | *CTL* | 4.3± 0.7 | - | - | - |
|  |  | *VaD* | 4.6± 0.9 | 0.8 | 0.51 | - |
|  | *Egr2* | *CTL* | 10.3± 2.6 | - | - | - |
|  |  | *VaD* | 11.4± 3.1 | 0.5 | 0.55 | - |
| *Cellular signaling* | *c-Jun* | *CTL* | 5.0 ± 0.5 | - | - | - |
|  |  | *VaD* | 4.7± 1.0 | 1.5 | 0.46 | - |
|  | *c-Fos* | *CTL* | 8.4± 1.7 | - | - | - |
|  |  | *VaD* | 6.3± 0.8 | 4.3 | 0.04* | opposite |
|  | *Notch1* | *CTL* | 8.6± 0.2 | - | - | - |
|  |  | *VaD* | 8.2± 0.3 | 1.4 | 0.04 | opposite |
| *Lipid metabolism* | *PLIN4* | *CTL* | 13.4± 1.1 | - | - | - |
|  |  | *VaD* | 12.9± 0.7 | 1.5 | 0.33 | - |
|  | *Lcn2* | *CTL* | 15.0± 2.0 | - | - | - |
|  |  | *VaD* | 12..3± 1.9 | 6.3 | 0.05 | consistent |
| *Infalmmatioon*  *Vascular remodeling* | *Klf4* | *CTL* | 13.9± 1.5 | - | - | - |
|  |  | *VaD* | 11.9± 1.5 | 4 | 0.05 | opposite - |
|  | *Cyp1b1* | *CTL* | 5.6± 1.2 | - | - | - |
|  |  | *VaD* | 5.8± 0.5 | 0.9 | 0.78 | - |
|  | *Angptl4* | *CTL* | 8.2± 0.8 | - | - | - |
|  |  | *VaD* | 6.9± 0.7 | 2.3 | 0.03 | consistent |
