## Supplementary material for "Systemic inflammation causes microglial dysfunction with a mixed AD-like pathology": Suppl. Fig. Legends

**Supplementary Figure Legends**

**Supplementary Figure 1. PolyI:C double immune challenge does not lead to infiltration of peripheral leukocytes into the brain**. Blood and brain specimens were collected at 3 months (3 weeks post second immune challenge) and at 6 months in PolyI:C (PP) & saline controls (NN). **(A)** Percentage of blood and brain neutrophils at 3 and 6 months in the PP and NN mice. **(B)** Total polymorphonuclear cells (PMN) frequency in the whole blood and in the brain. **(C)** Percentage of monocyte population in the blood and brain after the treatment. Percentage values are represented over % CD45+ cell frequency, data represented as mean ±SEM. n=4-6 mice per age group and treatment condition, *p<0.05 Welch’s t-test.

**Supplementary Figure 2. Visible tauopathy in the hippocampus:** The expression of p-tau in the hippocampal CA1 field in the PolyI:C and saline treated mice. **(A)** Immunolabeling for p-Tau in all age groups. **(B)** Analysis of the Mean fluorescence intensity (MFI) of pTau in the CA1 of 16 months PolyI:C and saline treated mice. Scale bar in A is 25 μm. *p<0.05 Student's t-test, n= 3 mice per condition.

**Supplementary Figure 3: Dynamic microglia soma morphological changes across PolyI:C aging mice:** **(A)** Distribution analysis showed that following PolyI:C treatment (PP) hippocampal microglial activity is associated with an increased cell soma area at3 months (KS2 test, Two Sample Kolmogorov-Smirnov Test (KS), D=0.25, P<0.005), subtle changes at 6 months and increased cell body area at 9 months (KS2 test, D=0.39, p<0.005) and back to normalcy in 16 months aged mice. **(B)** Increased soma perimeter was observed not in all age groups (3 months, KS2, D=0.24, p<0.005; 6 months, KS2, D=0.11, p=0.22; 9 months, KS2, D=0.28, p<0.005; 16 months, KS2, D=0.19, p=0.33. **(C)** Soma circularity showed no changes at 3 months; less circular or irregular at the intermediate stages (6 months, KS2, D=0.16, p<0.05; 9 months, KS2, D=0.28, p<0.005) until the 16 months aged mice where the shape appeared to shift more circular compared to microglia in the control animals (KS2, D=0.48, p<0.005). n=3 mice per age group and treatment.

**Supplementary Figure 4. SYNGO of DEGs**. Sunburst plot for Gene counts blasted on the SynGO database of DEGs in the aging PolyI:C mice as compared to saline controls at **(A)** 3months **(B)** 6months **(C)** 9 months **(D)** 16months. Color coding based on the gene count values indicated at each age.

**Supplementary Figure 5. Map of progressive changes reflected by functional interactomes.** Functional gene enrichment analysis for statistically most significant pathways of DEGs in the aging PolyI:C mice as compared to saline controls visualized in ClueGO software based on KEGG, Reactome, WikiPathways at **(A)** 3months **(B)** 6months **(C)** 9 months **(D)** 16months. Upregulated and downregulated genes are highlighted in blue and red text respectively. Each node represents the biological pathway and color of the node represents the functional group.
