## Supplementary figures and images for "Systemic inflammation causes microglial dysfunction with a mixed AD-like pathology"

### Suppl. Fig 1

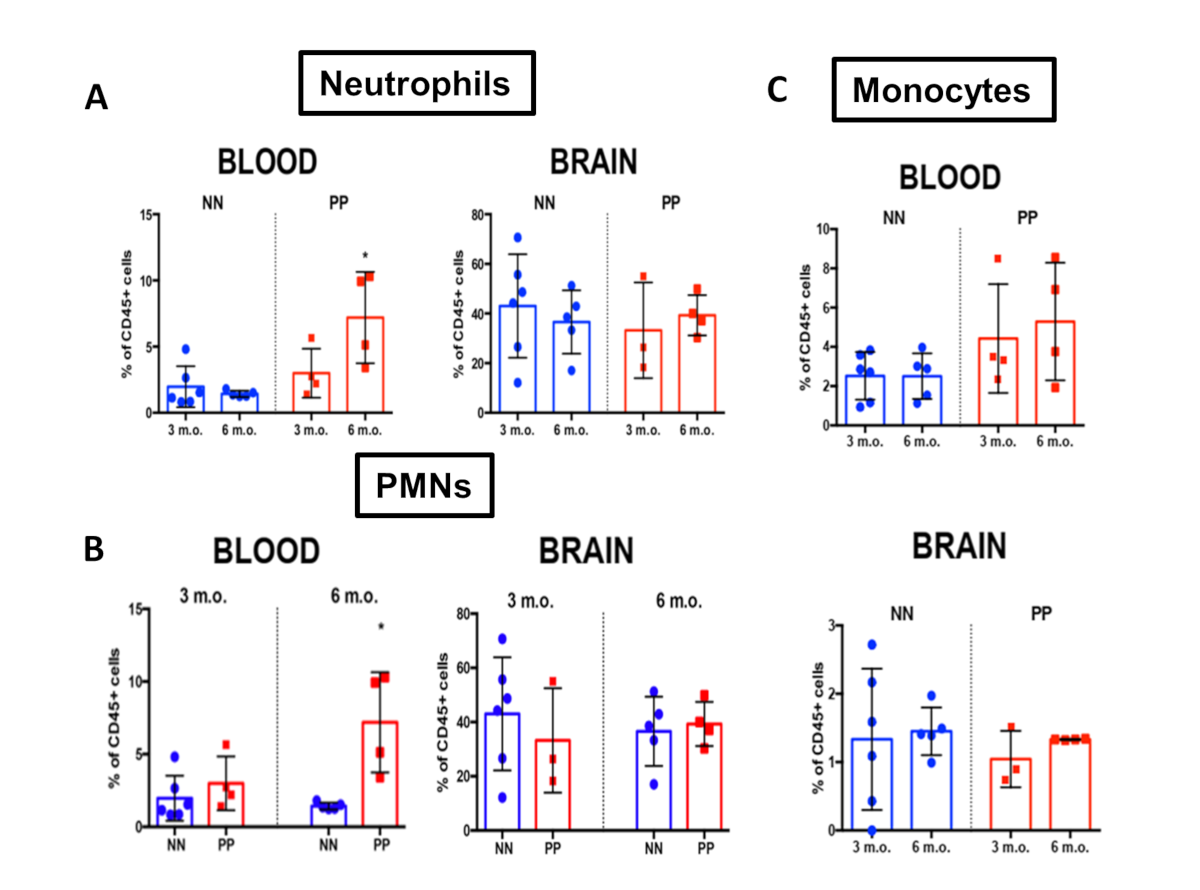

### Suppl. Fig 2

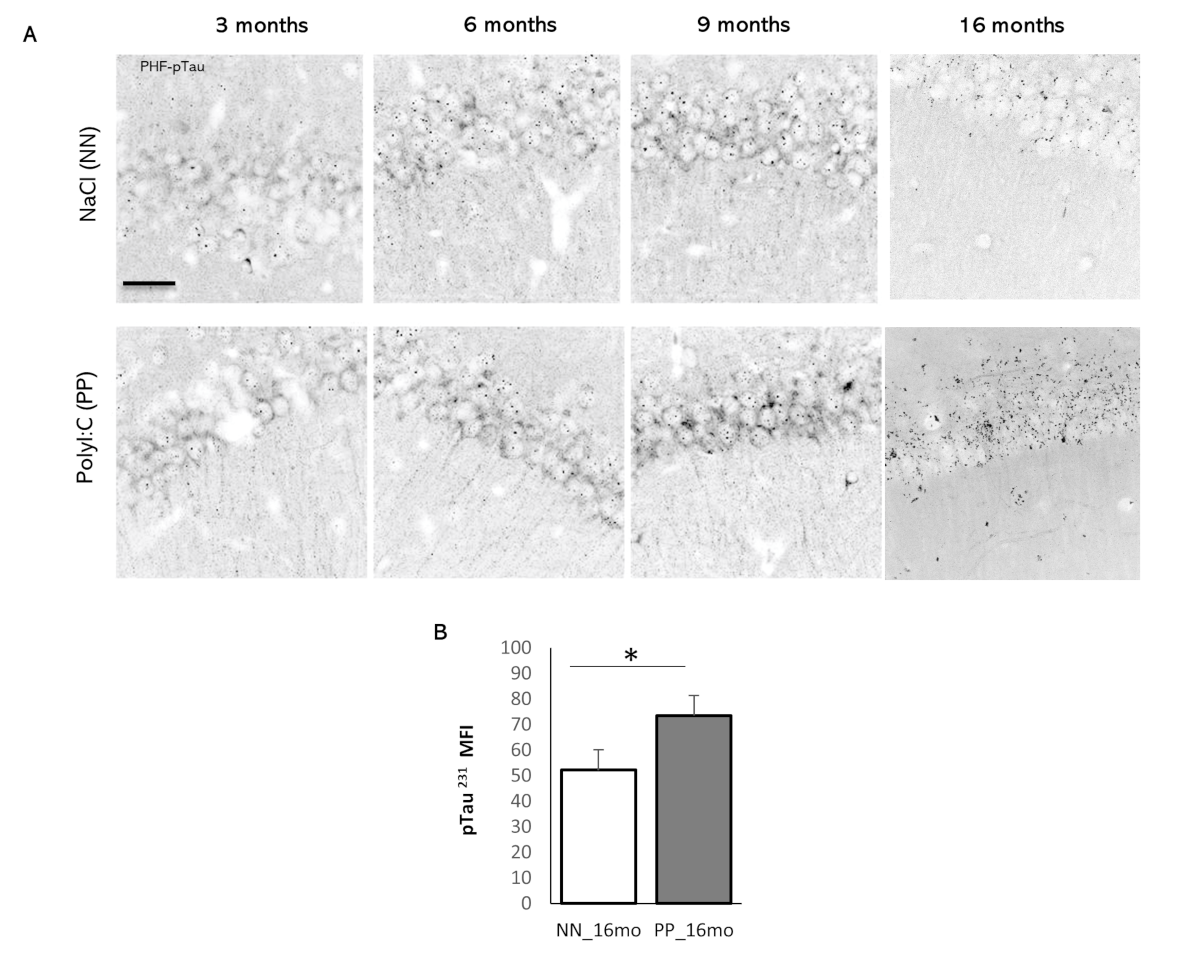

### Suppl. Fig 3

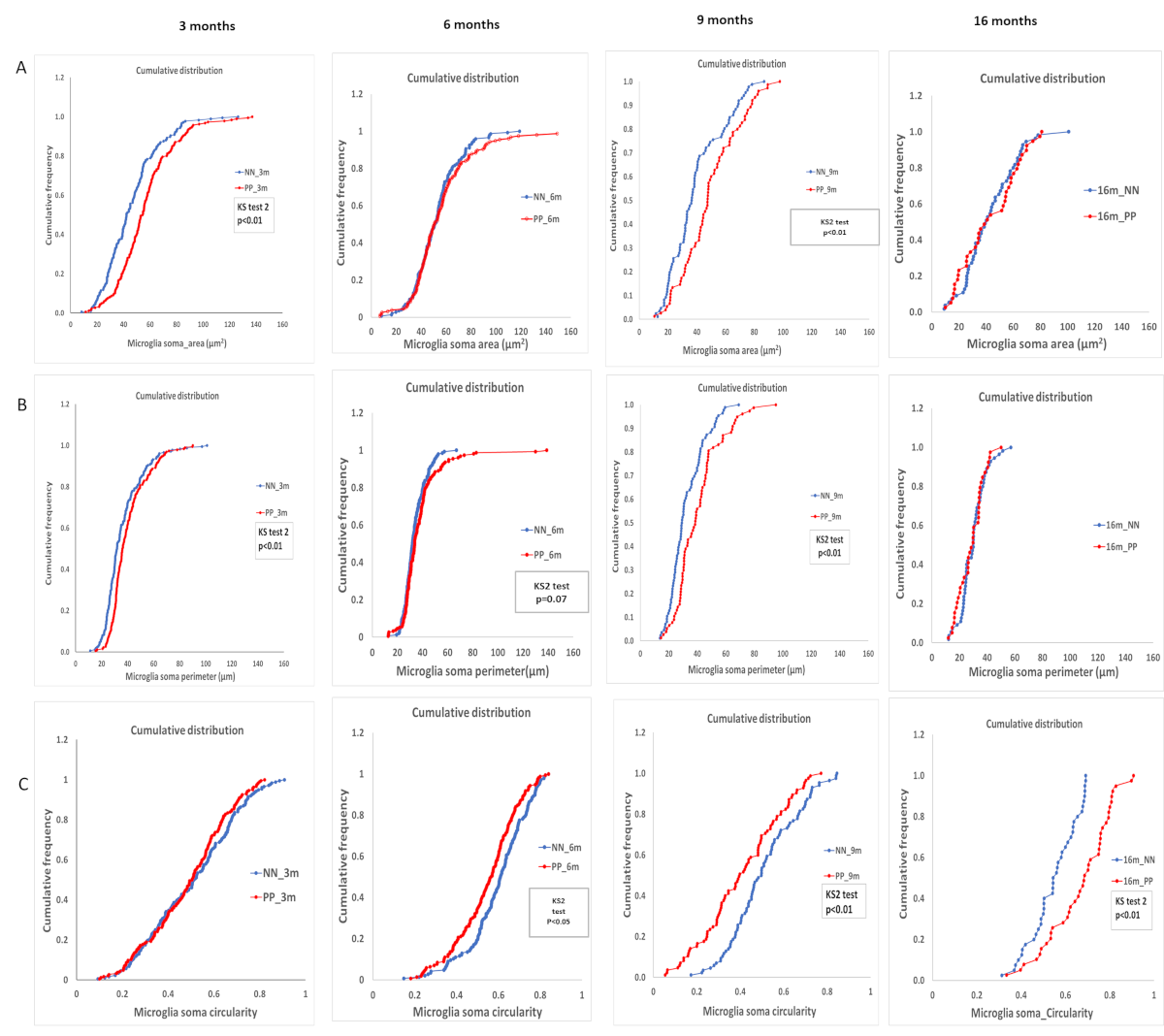

### Suppl. Fig 4

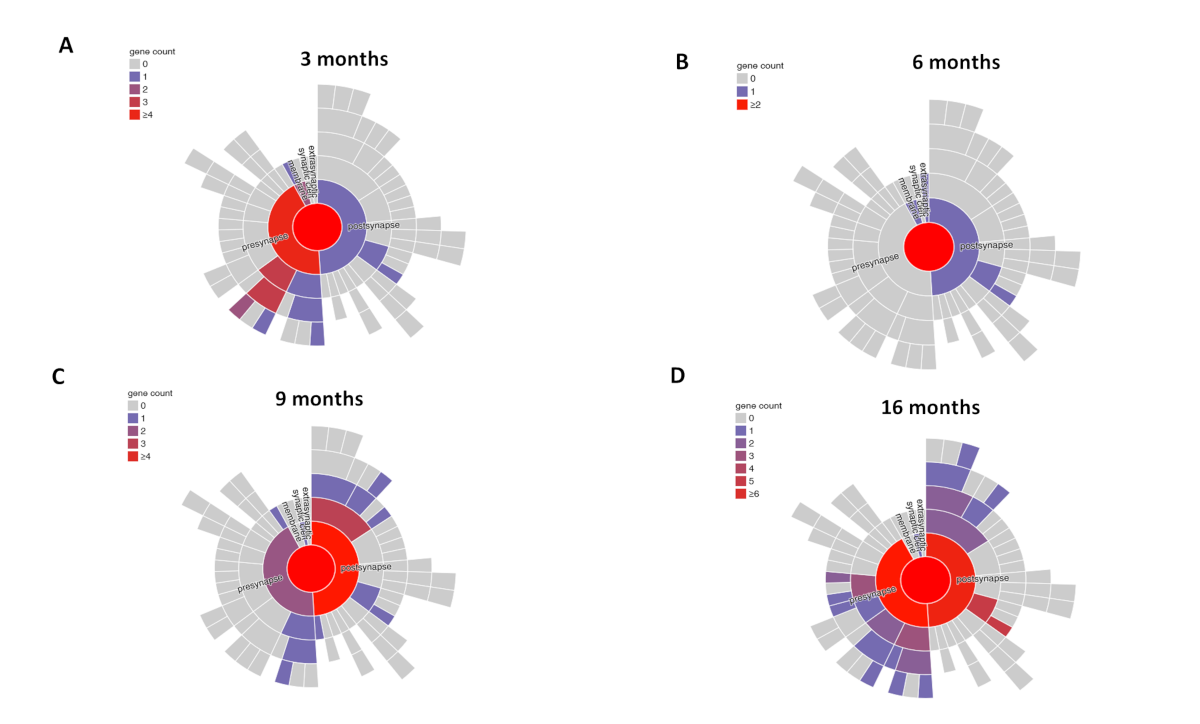

### Suppl. Fig 5

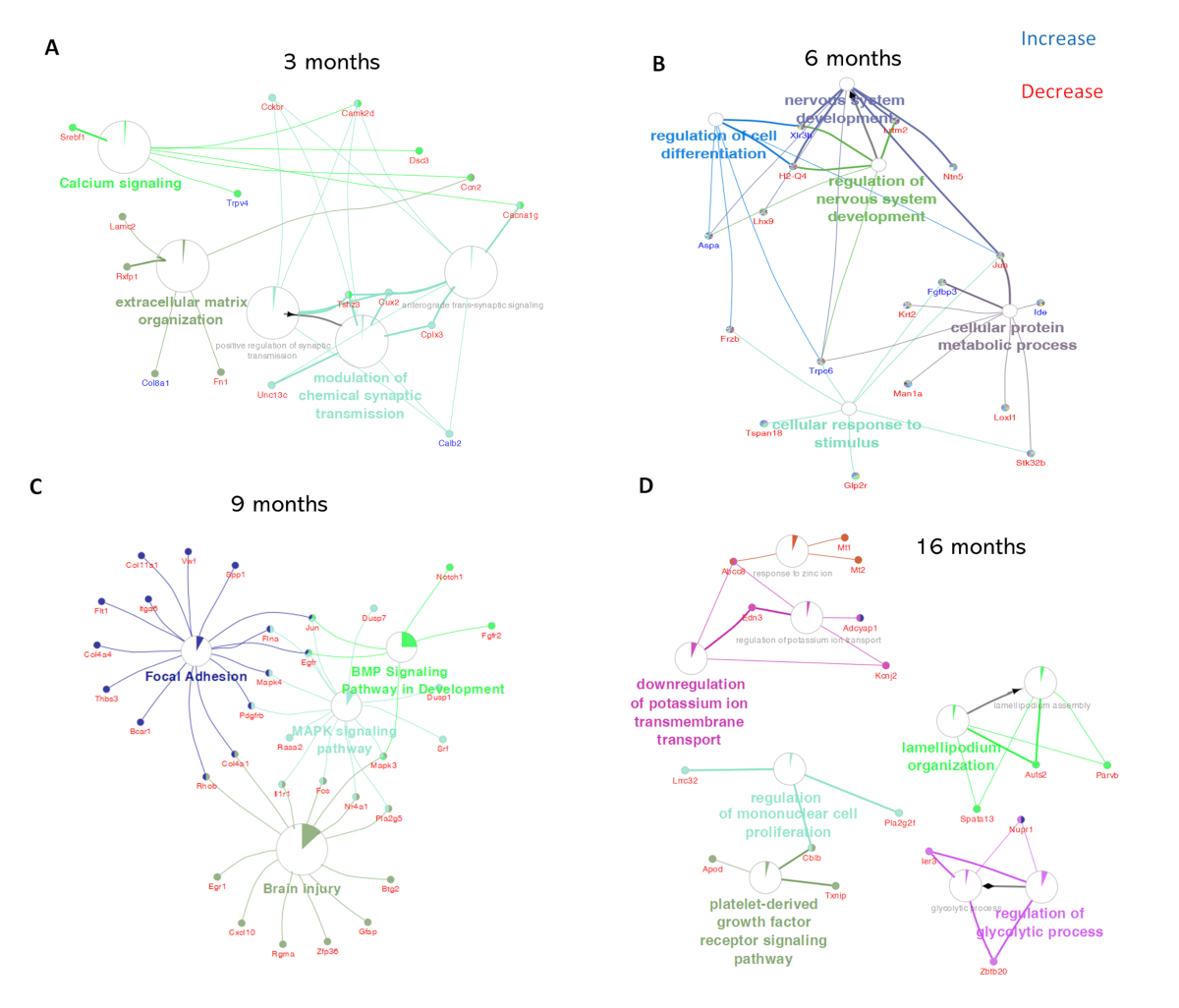
