## Supplementary material for "Systemic inflammation causes microglial dysfunction with a mixed AD-like pathology": Suppl. Material

**Supplementary Materials and Methods**

**ImageJ macro_ Y maze spatial alternation task**

*run("In");*

*makeOval(74, 45, 175, 172);*

*run("Crop");*

*run("In");*

*setAutoThreshold();*

*//run("Threshold...");*

*setThreshold(0, 30);*

*run("Threshold", "thresholded remaining black stack");*

*run("Despeckle", "stack");*

*run("Variance...", "radius=1 stack");*

*Arm A_makePolygon(34,24,43,18,76,52,66,61);*

*run("Plot Z-axis Profile");*

*Arm B_makePolygon(81,74,82,90,145,83,144,70);*

*run("Plot Z-axis Profile");*

*Arm C_makePolygon(74,91,84,94,59,160,44,151);*

*run("Plot Z-axis Profile");*

**ImageJ macro_ Elevated O maze**

*run("In");*

*run("In");*

*run("In");*

*makeOval(226, 120, 248, 248);*

*run("Crop");*

*run("In");*

*run("In");*

*run("In");*

*setAutoThreshold();*

*//run("Threshold...");*

*setThreshold(0, 30);*

*run("Threshold", "thresholded remaining black stack");*

*run("Despeckle", "stack");*

*run("Variance...", "radius=1 stack");*

*makePolygon(44,42,96,10,152,9,198,34,180,56,140,36,96,40,60,62);*

*run("Plot Z-axis Profile");*

*makePolygon(67,181,98,201,150,200,186,176,206,197,154,226,92,226,51,201);*

*run("Plot Z-axis Profile");*

**ImageJ macro_ Open field exploration**

*run("In");*

*run("In");*

*run("In");*

*makeOval(93, 165, 496, 162);*

*run("Crop");*

*run("In");*

*run("In");*

*run("In");*

*run("In");*

*setAutoThreshold();*

*//run("Threshold...");*

*setThreshold(0, 30);*

*run("Threshold", "thresholded remaining black stack");*

*run("Despeckle", "stack");*

*run("Variance...", "radius=1 stack");*

*makeOval(0, 0, 496, 160)*

*run("Plot Z-axis Profile");*

*makeOval(70, 40, 390, 92);*

*run("Plot Z-axis Profile");*

**ImageJ macro_ Light/dark test**

*run("In");*

*run("In");*

*run("In");*

*//setTool("rectangle");*

*makeRectangle(43, 59, 122, 92);*

*run("Crop");*

*run("Auto Threshold...", "method=[Try all]");*

*setAutoThreshold("Default");*

*//run("Threshold...");*

*setThreshold(0, 30);*

*run("Threshold", "thresholded remaining black stack");*

*run("Despeckle", "stack");*

*run("Plot Z-axis Profile");*

**ImageJ macro_Microglia Soma analysis**

*selectWindow("Image");*

*run("Z Project...", "projection=[Max Intensity]");*

*run("Set Scale...", "distance=8.8106 known=1 pixel=1 unit=micron global");*

*setOption("ScaleConversions", true);*

*run("8-bit");*

*run("Duplicate...", " ");*

*//run("Brightness/Contrast...");*

*run("Enhance Contrast", "saturated=0.35");*

*run("Apply LUT");*

*run("Gray Scale Attribute Filtering", "operation=Opening attribute=Area minimum=80 connectivity=8");*

*run("Morphological Filters", "operation=Opening element=Octagon radius=5");*

*setAutoThreshold("MaxEntropy");*

*//run("Threshold...");*

*setAutoThreshold("MaxEntropy");*

*run("ROI Manager...");*

*roiManager("Show All");*

*//setTool("wand");*

*roiManager("Add");*

*run("Set Measurements...", "area centroid center perimeter fit shape integrated area_fraction redirect=None decimal=5");*

*roiManager("Select", 1);*

*run("Select All");*

*roiManager("Select", newArray(0,1));*

*roiManager("Measure");*

**ImageJ macro_ Microglia skeleton analysis**

*outputFolder=getDirectory("C:\Users\ images\output");*

*dir =getDirectory("image");*

*name= getTitle();*

*path= outputFolder+name;*

*//run("Brightness/Contrast...");*

*run("Enhance Contrast", "saturated=0.35");*

*run("Apply LUT");*

*run("Unsharp Mask...", "radius=8 mask=0.60");*

*run("Despeckle");*

*setAutoThreshold("Default dark");*

*//run("Threshold...");*

*setAutoThreshold("Otsu dark");*

*//setThreshold(102, 255);*

*setOption("BlackBackground", false);*

*run("Convert to Mask");*

*run("Despeckle");*

*run("Close-");*

*run("Remove Outliers...", "radius=3 threshold=50 which=Bright");*

*run("Skeletonize (2D/3D)");*

*run("Analyze Skeleton (2D/3D)", "prune=none show display");*

*selectWindow("Tagged skeleton");*

*saveAs("tiff",path);*

*run("Close");*

**ImageJ macro_Amyloid analysis**

*run("Z Project...", "projection=[Max Intensity]");*

*run("8-bit");*

*run("Set Scale...", "distance=3.2055 known=1 pixel=1 unit=micron global");*

*setAutoThreshold("Moments dark");*

*//setThreshold(11, 255);*

*setOption("BlackBackground", false);*

*run("Convert to Mask");*

*run("Analyze Particles...", "size=5-Infinity show=Outlines display exclude include summarize");*

**ImageJ macro_Lcn2 analysis**

*run("Z Project...", "projection=[Max Intensity]");*

*run("8-bit");*

*run("Set Scale...", "distance=3.2056 known=1 pixel=1 unit=Âµm global");*

*//run("Brightness/Contrast...");*

*run("Enhance Contrast", "saturated=0.35");*

*run("Apply LUT");*

*run("Subtract Background...", "rolling=50");*

*run("Median...", "radius=0.1");*

*setAutoThreshold("Default dark");*

*//run("Threshold...");*

*setAutoThreshold("Otsu dark");*

*//setThreshold(58, 255);*

*setOption("BlackBackground", false);*

*run("Convert to Mask");*

*run("Set Measurements...", "area mean centroid perimeter integrated median area_fraction redirect=None decimal=3");*

*run("Analyze Particles...", "size=0.5-20.00 show=[Bare Outlines] display exclude include summarize");*
